# Model-based dimensionality reduction for single-cell RNA-seq using generalized bilinear models

**DOI:** 10.1101/2023.04.21.537881

**Authors:** Phillip B. Nicol, Jeffrey W. Miller

## Abstract

Dimensionality reduction is a critical step in the analysis of single-cell RNA-seq (scRNA-seq) data. The standard approach is to apply a transformation to the count matrix followed by principal components analysis (PCA). However, this approach can induce spurious heterogeneity and mask true biological variability. An alternative approach is to directly model the counts, but existing methods tend to be computationally intractable on large datasets and do not quantify uncertainty in the low-dimensional representation. To address these problems, we develop scGBM, a novel method for model-based dimensionality reduction of scRNA-seq data using a Poisson bilinear model. We introduce a fast estimation algorithm to fit the model using iteratively reweighted singular value decompositions, enabling the method to scale to datasets with millions of cells. Furthermore, scGBM quantifies the uncertainty in each cell’s latent position and leverages these uncertainties to assess the confidence associated with a given cell clustering. On real and simulated single-cell data, we find that scGBM produces low-dimensional embeddings that better capture relevant biological information while removing unwanted variation.

## 1 Introduction

Single-cell RNA sequencing (scRNA-seq) is a revolutionary technology that allows gene expression to be profiled at the level of individual cells (Saliba et al., 2014). This enables the identification of novel cell types that play critical roles in biological processes. However, the increased resolution provided by scRNA-seq comes at the cost of introducing several statistical and computational challenges. One major challenge is the large size of scRNA-seq datasets, which often contain millions of cells (Cao et al., 2019); thus, new methods must be computationally scalable (Lähnemann et al., 2020). Another challenge is that the extreme sparsity and discreteness of scRNA-seq count data make traditional statistical models based on normal distributions inappropriate (Vallejos et al., 2017). A third major challenge is that cell-level measurements are noisy and contain relatively little information compared to bulk sequencing, making it important to quantify the uncertainty in these measurements and propagate it to downstream analyses (Lähnemann et al., 2020).

Due to the large size of single-cell datasets, it is standard practice to use a dimensionality reduction technique such as principal components analysis (PCA) before clustering and other downstream analyses (Luecken and Theis, 2019). Researchers typically apply a transformation to the count matrix before running PCA, since applying PCA directly to the count matrix can introduce strong technical artifacts. Unfortunately, commonly used transformations such as log(1+*x*) can still lead to substantial biases in the subsequent PCA results (Townes et al., 2019). This has led to the current leading approach of transforming the counts using the Pearson residuals of a probabilistic model, as done by scTransform (Hafemeister and Satija, 2019). However, we find that even scTransform leads to undesirable biases that can mask true signals and upweight technical artifacts. Indeed, it seems unlikely that there exists a transformation that satisfies the necessary requirements simultaneously (Van Eeuwijk, 1995).

Alternatively, some methods perform dimensionality reduction directly using a probabilistic model of the count data matrix. These methods can avoid the artifactual biases of simple transformations and, further, can provide principled uncertainty quantification for downstream analyses and visualization. The GLM-PCA method of Townes et al. (2019) models the entries of the count matrix using a Poisson or negative-binomial distribution and estimates latent factors in the log space, however, GLM-PCA suffers from slow runtime and convergence issues on single-cell datasets with millions of cells (Lause et al., 2021). ZINB-WAVE (Risso et al., 2018) employs a similar model that assumes a zero-inflated negative-binomial distribution for the counts, but it has been noted that ZINB-WAVE can take days to run on large datasets, and furthermore, several studies have shown that the distribution of UMI counts is not zero inflated (Svensson, 2020; Sarkar and Stephens, 2021; Townes et al., 2019). This has prompted the development of the faster version, NewWave (Agostinis et al., 2022), however, we find that even NewWave is computationally burdensome on large datasets. The scVI method of Lopez et al. (2018) uses a variational autoencoder to learn a low-dimensional embedding, but a disadvantage is that this embedding may not be easily interpretable (Svensson et al., 2020), in contrast to PCA where the embedding is defined by linear combinations of genes.

An advantage of a model-based approach to dimensionality reduction is that one can, in principle, quantify uncertainty in the low-dimensional representation of cells. Understanding the confidence in each cell’s latent position is useful for discerning whether the clusters represent biologically distinct groups as opposed to technical or computational artifacts. However, to the best of our knowledge, none of the existing methods use uncertainty in the low-dimensional embedding to provide uncertainty quantification in downstream analyses.

In this paper, we introduce scGBM, a novel approach to dimensionality reduction for scRNA-seq data. Starting from the same underlying model as in GLM-PCA (Townes et al., 2019), we provide three key innovations. First, we develop a new estimation algorithm that is faster than existing approaches and scales up to datasets with millions of cells. Second, we quantify uncertainty in the low-dimensional embedding, enabling calibrated inference in downstream analyses. Third, we use these uncertainties to define a *cluster cohesion index* (CCI) that measures the stability of each cluster and the relationships between clusters. The CCI provides a quantitative tool for assessing which clusters represent biologically distinct populations, rather than simply being artifacts of sampling variability. Further, when analyzing numerous clusters, the CCIs reveal relationships that would be difficult to see with scatterplots. We evaluate the performance of scGBM on real and simulated data, finding that in many cases the current leading approaches are unable to capture true biological variability, while scGBM is successful in doing so.

The article is organized as follows. In Section 2, we demonstrate some of the limitations of current leading methods for scRNA-seq dimensionality reduction. In Section 3, we introduce our proposed methodology. Section 4 contains empirical results comparing the performance of our method versus leading approaches on simulated and real data. We conclude in Section 5. An scGBM R package is available for download.^1^

## 2 Motivation

It is known that fundamental issues can arise from the commonly used approach of running PCA on log(1+*x*) transformed scRNA-seq data. For instance, Townes et al. (2019) showed that the first principal component is strongly correlated with the number of zeros per cell, even on null data. Methods using count models have been developed to address these issues, however, we find that the current leading approaches still have significant limitations. One of the most popular methods is scTransform (Hafemeister and Satija, 2019), implemented in the widely used Seurat package (Satija et al., 2015). Let *Y* ∈ ℝ^*I×J*^ be the matrix of unique molecular identifier (UMI) counts, where *I* is the number of genes and *J* is the number of cells. The idea behind scTransform is to apply PCA to the matrix of Pearson residuals obtained by fitting negative-binomial GLMs to the count matrix *Y*. Specifically, for each row *i*, scTransform fits the following model to data *Y*_*i*1_, …, *Y*_*iJ*_ :

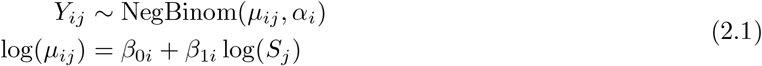

where 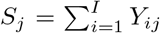 and NegBinom(*µ, α*) is the negative-binomial distribution with mean *μ*and variance *μ*+ *µ*^2^*/α*. Then, PCA is applied to the matrix *Z* = [*Z*_*ij*_] *∈* ℝ^*I×J*^ where

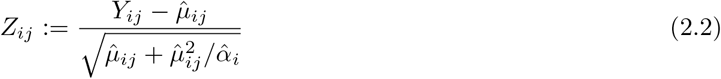

are the Pearson residuals and 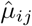 and 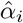 are regularized versions of the estimated parameters for the model in Equation 2.1. Lause et al. (2021) argued that the scTransform model (Equation 2.1) is overspecified since log(*S*_*i*_) can be interpreted as an offset which would imply that *β*_1*i*_ = 1. Additionally, Lause et al. (2021) showed that the overdispersion estimates 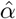 can have significant bias. We demonstrate potential shortcomings of scTransform using (i) a simulation with three cell types differentiated by single marker genes, and (ii) a simulation with two cell types differentiated by log-fold changes of equal magnitude across all genes.

### Single marker genes simulation

In simulation (i), we generated data for *J* = 1000 cells from three cell types, where cell types A and B are each distinguished by overexpressing a single marker gene. Specifically, relative to cell type C, cell type A overexpresses gene 1 and cell type B overexpresses gene 2, while the remaining 998 genes have identical mean expression across the three cell types (Figure 1a); see Section S3 for full details of the simulation. We applied scTransform to this dataset and found that, in terms of PCs 1 and 2, scTransform does not reveal the distinction between these three cell types (Figure 1b,c). The reason why this occurs is that scTransform implicitly standardizes the scale of the genes by dividing by the standard deviation, so all genes end up getting similar weight in the PCA. Additionally, the overdispersion parameter *α*_*i*_ absorbs some of the latent variability in the first two genes, further reducing the signal. This is illustrated in Figure 1d, which shows that the variance of the Pearson residuals from the 998 noise genes is comparable to (and, in fact, larger than) that of the two marker genes. Thus, the normalization procedure used by scTransform upweights the noise genes and downweights the signal genes.

**Figure 1.**
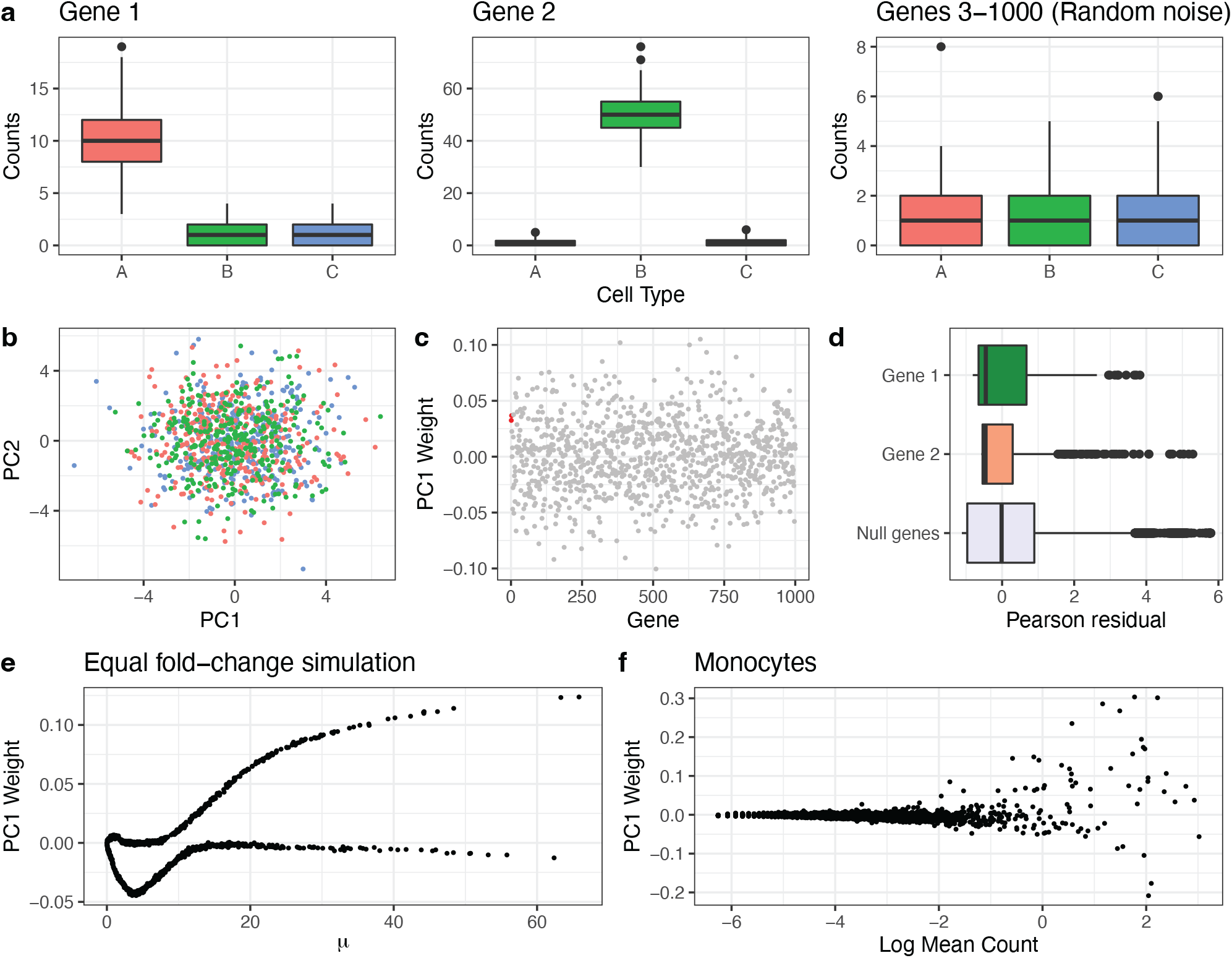
Limitations of scTransform. **a**. Single marker genes simulation: Boxplots of simulated counts for 1000 cells from 3 cell types. Gene 1 is overexpressed in cell type A and gene 2 is overexpressed in cell type B. The remaining genes are Poisson(1) random noise. **b**. The scores obtained from applying PCA to the Pearson residuals computed by scTransform. **c**. The corresponding weight of each gene in the first principal component. **d**. Boxplots of Pearson residuals for gene 1, gene 2, and the remaining “noise” genes. **e**. Equal log-fold change simulation: Two cell types such that all genes have an absolute log_2_ fold change of 1 between the two cell types. The weight given to each gene in the first principal component depends on the baseline expression *µ*. **f**. Purified monocytes data: Applying scTransform to monocytes from 10X genomics, the genes with the largest baseline mean (in terms of log mean expression) tend to have the largest PC weights.

### Equal log-fold change simulation

In simulation (ii), we generated *J* = 1000 cells from two equally sized groups. In the first group, we randomly generated the mean of each gene *i* = 1, …, *I* by sampling *µ*_*i*_ *∼* Exponential(0.1) independently. In the second group, the mean of gene *i* was randomly set to either 2*µ*_*i*_ or *µ*_*i*_*/*2, with probability 1*/*2, independently. In other words, all genes are differentially expressed by a multiplicative factor (fold change) of 2 between the two groups of cells. Each count was generated as independent Poisson with the corresponding mean. Applying scTransform (Figure 1e), we see that the PC1 weights are strongly dependent on the base mean *µ*_*i*_. This is undesirable because, in real data, some genes tend to have higher (or lower) measured expression simply because they are more (or less) abundant due to baseline cell activity or due to technical gene-specific effects unrelated to biology. Moreover, the PC weights are commonly used as a measure of a gene’s influence or importance on the corresponding component (Soumillon et al., 2014; Petropoulos et al., 2016; Roden et al., 2006).

The reason why the peculiar dependence in Figure 1e occurs is that scTransform decomposes latent effects additively in the mean rather than in the log of the mean. Although scTransform adjusts for gene-specific and cell-specific intercepts in log space via log(*µ*_*ij*_) = *β*_0*i*_ + *β*_1*i*_ log(*S*_*j*_), the latent effects are decomposed in linear space because applying PCA to the Pearson residual matrix *Z* is equivalent to finding an approximate low-rank factorization *U ΣV* ^*T*^ *≈ Z* where *U* ∈ ℝ^*I×M*^, *V* ∈ ℝ ^*J×M*^, and Σ = diag(*σ*_1_, …, *σ*_*M*_), that is,

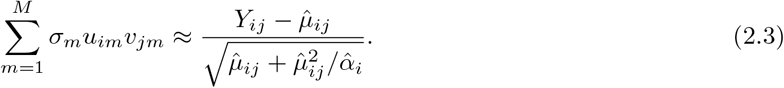

### Purified monocytes data

To illustrate this issue on real data, we considered a dataset of *J* = 2,612 purified monocytes downloaded from the 10X genomics website (www.10xgenomics.com). Since these data come from a single cell type, we expect minimal true latent variation. Since the true mean is unknown, we computed the log mean count for each gene and plotted it against the PC1 weights obtained from scTransform (Figure 1f). Although the weights are more variable than in the simulated example, we again observe that the genes with the largest baseline mean expression tend to have the largest weights.

## 3 Methods

In this section, we introduce our proposed model (Section 3.1), iteratively reweighted singular value decomposition (Section 3.2), step-by-step estimation algorithm (Section 3.3), projection technique for scaling to large matrices (Section 3.4), and uncertainty quantification approach (Section 3.5).

### 3.1 scGBM fits a Poisson bilinear model to the count matrix

Our proposed method, referred to as scGBM, handles normalization, transformation, and latent factorization in a unified way using a Poisson bilinear model. Specifically, we model the entries of the UMI count matrix *Y ∈* ℝ^*I×J*^ (with genes on rows and cells on columns) as

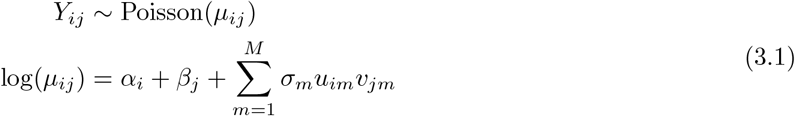

for *i* = 1, …, *I* and *j* = 1, …, *J*, where *α*_*i*_ is a gene-specific intercept, *β*_*j*_ is a cell-specific intercept, *U* := [*u*_*im*_] ∈ℝ^*I×M*^ is a matrix of latent factor loadings or weights, *V* := [*v*_*jm*_] ∈ ℝ ^*J×M*^ is a matrix of latent factor scores, and Σ := diag(*σ*_1_, …, *σ*_*M*_) *∈ ℝ*^*M ×M*^ is a diagonal matrix of scaling factors (or singular values) *σ*_*m*_ *>* 0. For intuition, this model can roughly be thought of as applying PCA inside the link function of a generalized linear model (GLM). As in PCA, we constrain the columns of *U* to be orthonormal and order the columns such that *σ*_1_ > … > *σ*_*M*_ *>* 0, so the first factor captures the greatest amount of latent structure. Identifiability constraints and more background on the model are provided in Section S1.1.

Following Choulakian (1996), we refer to the model in Equation 3.1 as a *generalized bilinear model* (GBM), but other names such as GAMMI (Van Eeuwijk, 1995) or eSVD (Lin et al., 2021) have been used in the statistical literature. We refer to Miller and Carter (2020) for an extensive discussion of the literature on GBMs. In the single-cell literature, Equation 3.1 is typically referred to as GLM-PCA (Townes et al., 2019). Although scGBM is based on the same model as GLM-PCA, there are some key differences between the two methods. First, scGBM is able to estimate cell-specific intercepts *β*_*j*_, whereas GLM-PCA treats the *β*_*j*_’s as fixed offsets. Second, GLM-PCA does not enforce identifiability constraints in the intercepts or the factors. Finally, GLM-PCA does not quantify uncertainty in the parameter estimates.

A GBM that only includes intercepts and latent factors, as in Equation 3.1, is likely to be sufficient for most single-cell studies. However, in some cases, it is beneficial to control for known batches such as groups of samples sequenced in different locations or on different dates. To achieve this, scGBM can estimate a batch-specific intercept for each gene; see Section S1.5 for details. More generally, the model can be augmented to include arbitrary sample covariates and gene covariates, as well as interactions between these (Miller and Carter, 2020). However, in this paper, we only consider binary sample covariates since other types of covariates are rarely used during this stage of single-cell analyses.

In contrast to many bulk RNA-seq models, the scGBM model assumes a Poisson outcome and thus does not allow for overdispersion. In fact, there is increasing evidence that the technical sampling distribution of UMI counts is close to Poisson and that any additional dispersion is due to biological variability such as heterogeneous cell types or cell states (Sarkar and Stephens, 2021; Lause et al., 2021). Since the goal of scGBM is to remove technical variability while preserving all biological variability for downstream analyses, it is natural to use a Poisson outcome distribution.

### 3.2 Fast estimation using iteratively reweighted singular value decomposition

Existing methods for fitting GBMs are not scalable to the large datasets encountered in scRNA-seq. To address this limitation, we propose a new approach to fitting the Poisson GBM that combines iteratively reweighted least squares (IRLS) and singular value decomposition (SVD). This pairing is natural since IRLS is the standard way to fit GLMs and SVD is the standard way to perform PCA.

Define *X* = *U*Σ*V* ^*T*^, and suppose we have current estimates 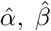 and 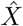. By a second-order Taylor approximation at these estimates (see Section S1.2 for details), maximizing the log-likelihood of the scGBM model in Equation 3.1 is approximately equivalent to solving the weighted low-rank problem

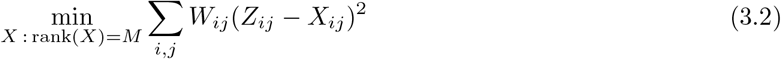

where 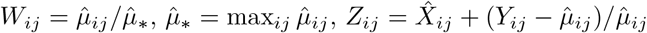, and 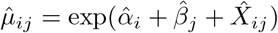. Unlike the unweighted case, there is not a closed-form solution to Equation 3.2, unfortunately. However, Srebro and Jaakkola (2003) introduced a simple and efficient method for solving this type of weighted low-rank problem (with fixed *W* and *Z*) by iteratively applying the equation

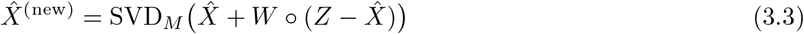

where SVD_*M*_ (*·*) denotes the truncated SVD of rank *M* and ∘ is the Hadamard product. In turn, when 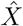 is held fixed, the maximum likelihood estimates of 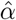 and 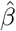 have closed-form solutions.

Thus, we alternate between updating 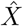 by applying Equation 3.3 with 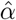 and 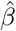 fixed, and updating 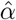 and 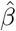 to their maximum likelihood estimates with 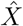 fixed. The truncated SVD automatically factors 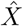 as 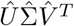, yielding estimates of the loadings, scores, and singular values. See Section 3.3 for a step-by-step description of the proposed algorithm, which we call *iteratively reweighted singular value decomposition* (IRSVD).

There are several advantages of the IRSVD algorithm. First, it is asymptotically faster than Fisher scoring, the technique used by Miller and Carter (2020) and Townes et al. (2019); see Section 3.3 for the computational complexity. Further, one can leverage special properties of Poisson GLMs to obtain vectorized updates for the intercepts—that is, the entries of *α* and *β* can be updated simultaneously—which greatly reduces the runtime in practice. Finally, the identifiability constraints such as orthogonality are preserved at every iteration of the algorithm.

### 3.3 Estimation algorithm and computational complexity

Our proposed IRSVD estimation algorithm for the scGBM model in Equation 3.1 is as follows.

- Initialize. Set 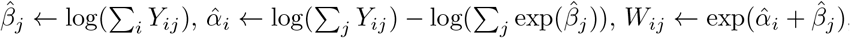 and

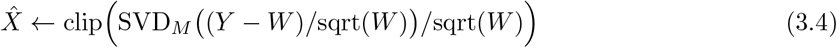

where square roots, division, and clip(*x*) = max(min(*x, c*), *−c*) are applied entry-wise, with *c* = 8.
- Iterate the following steps until convergence.

#### 1. Update intercepts

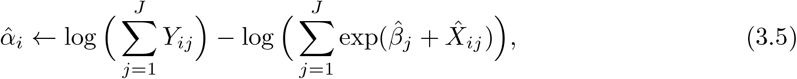

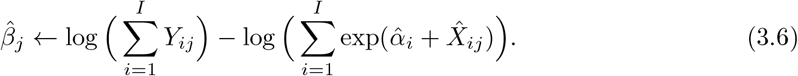

#### 2. Update latent factors

Compute 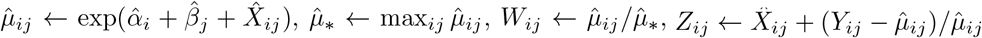, and

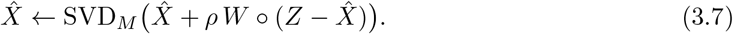

##### Output

Return the estimates 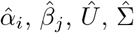, and 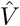 such that 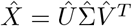.

For the convergence criterion, we terminate the algorithm when the relative change in log-likelihood is below a specified tolerance (default 10^*−*4^) or after a specified maximum number of iterations (default 100). As a default, we set *M* = 20. We find that a step size of *ρ* = 1 works well, but that the rate of convergence can be improved by using an adaptive scheme where *ρ* is increased by a multiplicative factor of 1.05 when the log-likelihood increases and is decreased by a multiplicative factor of 1*/*2 when the log-likelihood decreases. In addition, the latent factors update (Equation 3.7) can be modified to use Nesterov acceleration (Tuzhilina and Hastie, 2021); see Equation S1.5. Therefore, we use a slightly refined version of the algorithm employing Nesterov acceleration and the adaptive step size described above.

See Section S1.2 for the rationale behind each part of the algorithm. The initialization is based on solving the initial weighted low-rank problem exactly when *W* is rank 1. The intercept update step is based on maximum likelihood estimation holding 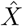fixed. In the latent factors update step, we augment the Srebro and Jaakkola (2003) iteration by allowing a step size *ρ >* 0 other than 1, and allowing *W* and *Z* to depend on the current value of 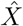 in order to form a local approximation to the weighted low-rank problem.

##### Computational complexity analysis

- **Initialize**. The truncated SVD of rank *M* takes *O*(*IJM*) time (Halko et al., 2011). The other initialization operations only take *O*(*IJ*) time, so this step is *O*(*IJM*) altogether.
- **Update intercepts**. Each 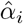 update takes *O*(*J*) time, and each 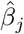 update takes *O*(*I*) time. Thus, the entire step takes *O*(*IJ*) time per iteration.
- **Update latent factors**. The entry-wise operations are *O*(*IJ*) and the truncated SVD of rank *M* takes *O*(*IJM*) time (Halko et al., 2011). Thus, this step takes *O*(*IJM*) time per iteration.

Hence, the runtime for initialization and each iteration of IRSVD is *O*(*IJM*). This is faster than the Fisher scoring algorithm of Miller and Carter (2020), which is *O*(*IJM* ^2^) per iteration. The quadratic complexity in *M* can be practically significant since *M* is often chosen to be around 20-50. Further, IRSVD is simpler than the Miller and Carter (2020) algorithm, making it easier to implement and yielding a smaller constant.

### 3.4 Scaling up by fitting on a subset of samples and projecting

Even with the improvement due to IRSVD, the computation time scales linearly with the number of samples, making it burdensome on datasets with millions of cells. Thus, to further improve scalability, we propose using a combination of subsampling and projection. Specifically, we first randomly select a subset of cells and run the algorithm in Section 3.3 to estimate the parameters using only the data from these cells. Then, holding *U* and *α* fixed at these estimated values, column *j* of the Poisson bilinear model in Equation 3.1 is simply a GLM with covariate matrix *U* and coefficients *β*_*j*_, *σ*_1_*v*_*j*1_, …, *σ*_*M*_ *v*_*jM*_, which can be fit using standard software. We refer to this as the “projection method”.

We refer to this subsampling-projection version of the algorithm as “scGBM-proj”, in contrast to running the algorithm on the full dataset, which we refer to as “scGBM-full”. The observation that fixing *U* or *V* yields a standard GLM has been used in previous RNA-seq techniques (Leek and Storey, 2007; Sun et al., 2012; Risso et al., 2014). Importantly, the GLMs can be fit in parallel since each column is processed independently. This also reduces memory requirements since the count matrix *Y* can remain stored in sparse form and only needs to be loaded into memory one column at a time, making it suitable for on-disk processing tools like *DelayedArray* (Pagès, 2020).

### 3.5 Quantifying uncertainty in the latent factors

Since scGBM performs dimensionality reduction using a statistical model, uncertainty in the low-dimensional representation can be quantified using the classical method of inverting the Fisher information matrix. Miller and Carter (2020) use this approach, accounting for joint uncertainty in *U* and *V* as well as their identifiability constraints by inverting the constraint-augmented Fisher information. However, this is computationally expensive on single-cell datasets.

Thus, to quantify uncertainty in *V*, we use a rough approximation by inverting the diagonal blocks of the Fisher information for *V*, that is, the submatrices *F*_1_, …, *F*_*J*_ ∈ ℝ ^*M ×M*^ in which entry (*m, m*^*′*^) of *F*_*j*_ is

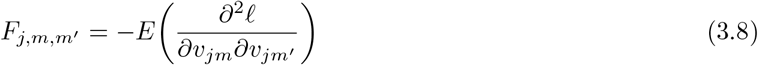

for *j* = 1, …, *J* and *m, m*^*′*^ = 1, …, *M*. For the Poisson bilinear model in Equation 3.1,

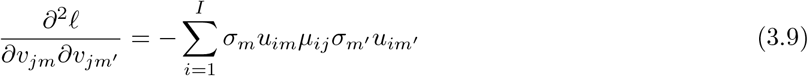

by differentiating the log-likelihood *ℓ*. Therefore, in matrix form,

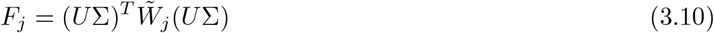

where 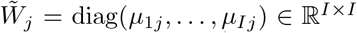. By plugging in parameter estimates and taking the square root of the diagonal entries of 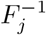, we obtain approximate standard errors for the estimates of *v*_*j*1_, …, *v*_*jM*_. This approximation will tend to underestimate the uncertainty in *V* since it treats the other parameters *α, β, U*, and Σ as fixed. Nonetheless, on simulated data, we find that it yields confidence intervals with coverage only slightly below the target coverage, for large *I* and *J* (Figure S1). Standard errors for *U* can be approximated in a complementary way.

## 4 Results

We evaluate the performance of scGBM by comparing to GLM-PCA (Section 4.1), testing the accuracy and speed of the projection method (Section 4.2), comparing to PCA coupled with scTransform or log(1 + *x*) transformation (Section 4.3), and applying our uncertainty quantification method to clustering (Section 4.4). For a description of all datasets used in this section, see Section S2.

### 4.1 Comparing to GLM-PCA

In this section, we compare the performance of our estimation algorithm (scGBM-full) to three algorithms provided by the GLM-PCA R package as implemented by Townes and Street (2020): Fisher scoring, Avagrad (Savarese et al., 2021), and stochastic gradient descent (SGD). We generated simulated datasets with *I* =

1000, *J* ∈ {2000, 10^5^}, and *M* = 10 by sampling *U* and *V* uniformly from the Stiefel manifold, setting Σ = diag(*σ*_1_, …, *σ*_*M*_) where 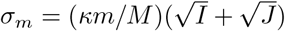, generating intercepts *α*_*i*_ and *β*_*j*_ independently from *N*(0, 1), and generating *Y* according Equation 3.1. On each dataset, we ran the estimation algorithms to produce parameter estimates Û 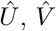, and 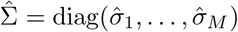.

To assess accuracy, we simulated 100 datasets as above with *J* = 2000 and *κ* = 2. For each *m* = 1, …, 10, Figure 2a shows the absolute value of the correlation between column *m* of (i) *U* and *Û* and (ii) *V* and 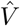. Additionally, for each dataset, we computed the root mean squared error between the true *X* = *U*Σ*V* ^*T*^ and the estimate 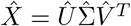 produced by each algorithm; see Figure 2b. In terms of both accuracy metrics, we found that scGBM-full performed significantly better (higher correlation, lower error) than GLM-PCA.

**Figure 2.**
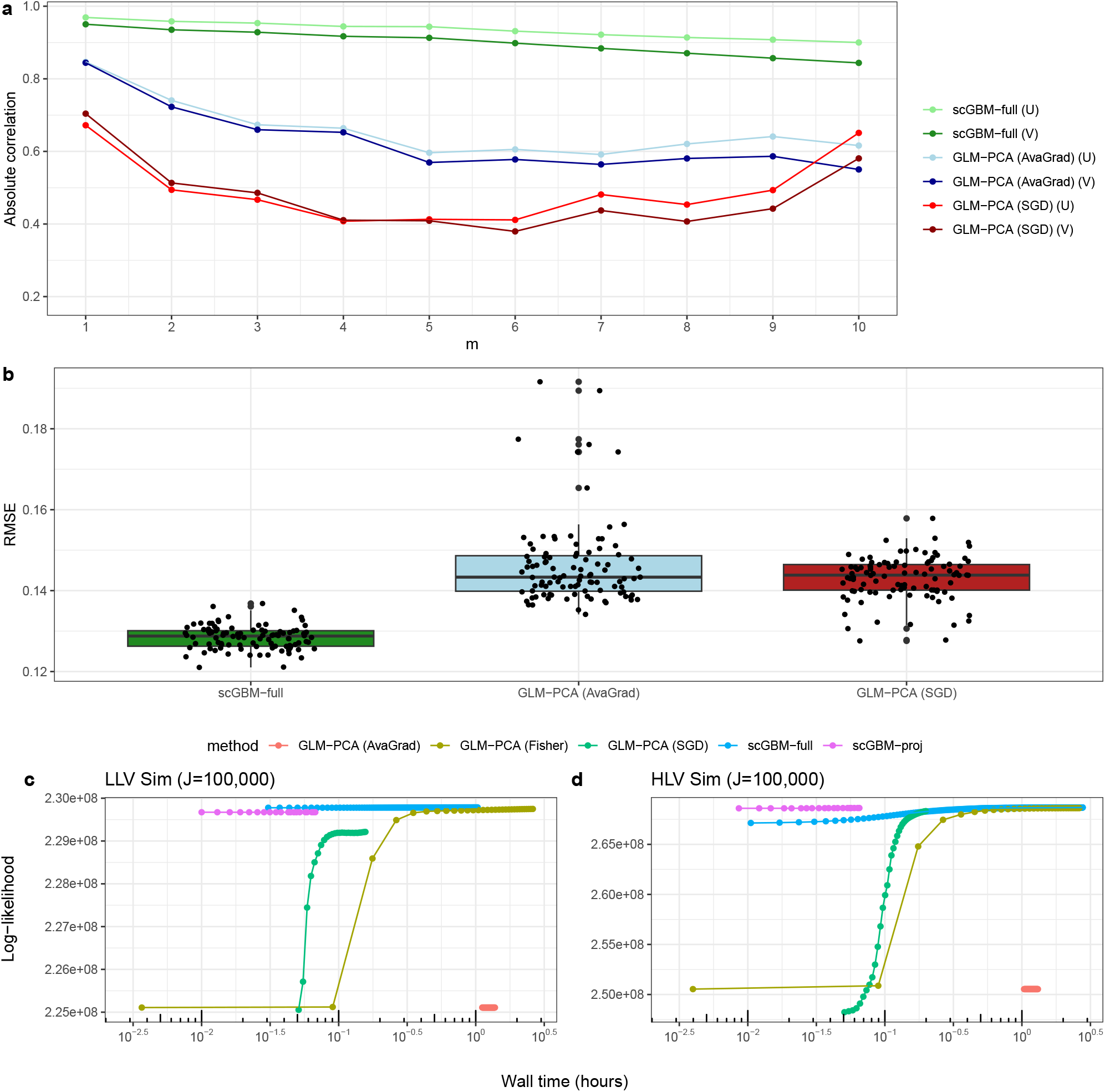
Comparison of scGBM-full (IRSVD) and GLM-PCA on simulated data. **a**. Absolute value of the correlation between estimated columns of *U* and *V* and the ground truth. Points shown are the median over 100 simulated datasets with *I* = 1000, *J* = 2000, and *M* = 10. **b**. Root mean squared error (RMSE) between the ground truth latent effects *X* = *U*Σ*V* ^*T*^ and the estimate 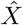 from each algorithm, for the 100 simulated datasets. GLM-PCA (Fisher) was not included in plots **a** and **b** since it has significantly longer runtime, making it computationally burdensome. **c**. Log-likelihood versus wall clock time on a low latent variability (LLV) simulated dataset with *I* = 1000 and *J* = 10^5^. **d**. The same plot for a high latent variability (HLV) simulated dataset with *I* = 1000 and *J* = 10^5^.

To compare computation time in a way that accounts for convergence rate and goodness of fit, we plot the log-likelihood Σ_*i,j*_ (*Y*_*ij*_ log(*µ*_*ij*_) *− µ*_*ij*_) (with additive constants removed) versus the wall clock time, that is, the elapsed time since the start of the algorithm. Figure 2 shows the results for datasets with *J* = 10^5^ under two simulation settings: *κ* = 2, representing low latent variability (Figure 2c), and *κ* = 5, representing high latent variability (Figure 2d). We observe that the initialization used by scGBM-full has significantly higher log-likelihood, and scGBM-full reaches better solutions more quickly than the GLM-PCA algorithms. In particular, GLM-PCA (AvaGrad) suffers from low log-likelihood and requires many runs to find a suitable learning rate.

For a real data comparison, Table 1 shows the runtime of scGBM-full and the three GLM-PCA algorithms on three real datasets: 10X immune cells (*J* = 3,994, Zheng et al., 2017), COVID-19 Atlas (*J* = 44,721, Wilk et al., 2020), and 10X mouse brain (*J* = 1,308,421, Lun and Morgan, 2020); see Table 1 caption for details. Figure S2 shows plots of log-likelihood versus runtime for the 10X immune cells and COVID-19 Atlas.

**Table 1:** Runtime comparison of scGBM, GLM-PCA, and NewWave on three datasets: 10X immune cells, COVID-19 Atlas, and 10X mouse brain; see Section S2 for dataset references. For each dataset, *I* = 1000 variable genes were selected using the Seurat package (Satija et al., 2015). Each method was run for a maximum of 100 iterations. For scGBM-proj, we used subsamples of sizes 400, 4,000, and 100,000 on the three datasets, respectively, and 64 cores were allocated for parallel processing. For GLM-PCA (SGD), the minibatch size was set equal to the subsample size used by scGBM-proj. For GLM-PCA, the learning rate (for Avagrad and SGD) and the penalty (for Fisher scoring) were set sufficiently small (or large) to prevent divergence. Latent factor dimensionality of *M* = 20 was used for all algorithms. When applying NewWave to the 10X brain data, the job timed out after 2907 minutes. *The runtime for GLM-PCA (Fisher) on the 10X mouse brain data was estimated by extrapolating the time taken for 10 iterations.

| Dataset | # of cells | scGBM-proj | scGBM-full | GLM-PCA (Fisher) | GLM-PCA (AvaGrad) | GLM-PCA (SGD) | NewWave |
| --- | --- | --- | --- | --- | --- | --- | --- |
| 10X immune cells | 3,994 | 0.14 min | 1.68 min | 55.72 min | 1.58 min | 1.72 min | 2.95 min |
| COVID-19 Atlas | 44,721 | 1.89 min | 22.77 min | 468.24 min | 19.52 min | 20.66 min | 35.31 min |
| 10X mouse brain | 1,308,421 | 57.67 min | 782.40 min | 9878.20 min* | 811.09 min | 505.82 min | >2907 min |

Overall, we found that scGBM-full was roughly an order of magnitude faster than GLM-PCA (Fisher). GLM-PCA (Avagrad) took approximately the same time as scGBM-full, and GLM-PCA (SGD) was similar or somewhat faster, however, GLM-PCA (Avagrad) and GLM-PCA (SGD) often converged to significantly less accurate solutions than scGBM-full (Figure 2a,b). Another limitation of the gradient-based methods implemented in GLM-PCA is that they are very sensitive to the learning rate. For instance, Table S1 shows an example where changing the learning rate by just 0.01 leads to divergence. This significantly increases the computational burden in practice, since the algorithm must search for a suitable learning rate.

### 4.1 Testing the performance of the projection method

Here, we assess the accuracy and computational efficiency of the projection method for scaling to large data (Section 3.4). To test it on real data, we consider the 10X immune cell and COVID-19 Atlas datasets from Table 1. We first applied scGBM-full to the whole datasets, obtaining 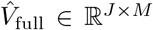, and then we used scGBM-proj with various subset sizes to obtain 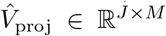. To quantify the accuracy of the projection method, we computed the absolute value of the correlation between 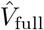 and 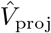 for each column *m* = 1, …, *M* ; see Figure 3. We found that the first column of *V* (that is, the one corresponding to the largest scaling factor, *σ*_1_) can be reliably estimated even using a small fraction of the cells to estimate *U*. These results suggest that using the projection method with *≈* 10-15% of the data is sufficient for capturing the most dominant sources of biological variation. We obtained similar results when testing the projection method on simulations with known ground truth; see Figure S3.

**Figure 3.**
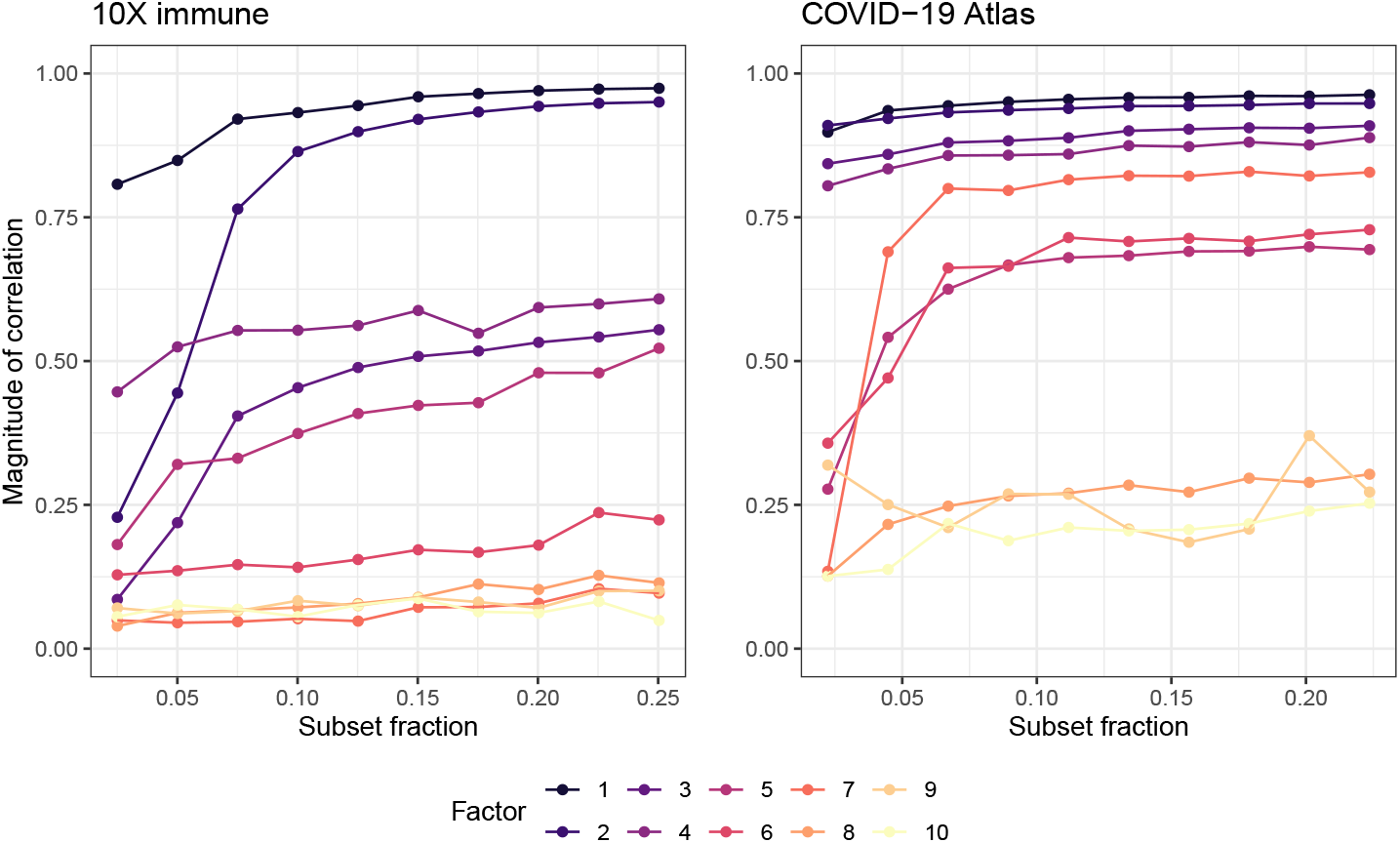
Testing the accuracy of the projection method. We first applied scGBM-full to the 10X immune cell dataset (*J* = 3,994) to estimate *V*. Then for subsets of various sizes, we used scGBM-proj to estimate *V*, and for each *m* = 1, …, 10, computed the correlation between the two estimates (scGBM-full and scGBM-proj) of the *m*th column of *V*. The points shown represent the median of the absolute value of the correlation, over 100 runs of scGBM-proj. As the subset fraction increases, scGBM-proj agrees more closely with scGBM-full, with 10-15% being sufficient to accurately capture the largest factors. Running the same analysis on the COVID-19 Atlas dataset (*J* = 44,721) yielded similar conclusions. The 10X mouse brain dataset was omitted due to the computational burden of numerous runs required.

To assess the computational efficiency of the projection method, Table 1 reports the runtime of scGBM-proj on the 10X immune cell, COVID-19 Atlas, and 10X mouse brain datasets. We find that scGBM-proj has significantly faster runtimes compared to the other algorithms. In particular, the total runtime on a dataset with 1.3 million cells was less than one hour using a high-performance computing environment with 64 cores. Figures 2c,d and S2 show the log-likelihood attained by scGBM-proj versus wall clock time on simulated and real data. These plots show that scGBM-proj consistently attains the highest log-likelihood for small wall clock times. On the data with high latent variation (HLV), scGBM-proj and scGBM-full converge to solutions with similar log-likelihood, but scGBM-proj does so much more rapidly.

### 4.3 Comparing to PCA coupled with scTransform or log(1 + *x*) transformation

Next, we assess the quality of scGBM results compared to two commonly used approaches, finding that it overcomes the limitations observed in Section 2. The first approach is “log+scale+PCA”, which applies PCA to the matrix log(1 +CPT) with all rows standardized to have mean 0 and standard deviation 1. Here, CPT denotes the *I × J* matrix of “counts per ten-thousand.” The second approach is SCT+PCA, which applies PCA to the matrix of Pearson residuals produced by scTransform as described in Section 2. Both methods are implemented in the *Seurat* package (Satija et al., 2015).

One might think that SCT+PCA and scGBM would yield very similar results since, as stated by Townes et al. (2019), applying PCA to the Pearson residuals can be viewed as an approximation to fitting the Poisson bilinear model used by scGBM. This approximation is justifiable when the scale of the latent variation is small (see the discussion following Proposition S1.1), however, when the scale of the latent variation is large, SCT+PCA and scGBM can produce very different results.

#### Single marker genes simulation

To illustrate, first consider the single marker genes simulation from Figure 1. Figure 4a plot the factor scores from the three methods, showing that scGBM is the only method able to distinguish the three simulated cell types. Even after applying tSNE (Van der Maaten and Hinton, 2008) to SCT+PCA and log+scale+PCA, there is no separation between the cell types (Figure S4). This indicates that the biological signal is not sufficiently captured by any of the PC dimensions in SCT+PCA or log+scale+PCA. In terms of the first factor loading, scGBM clearly prioritizes genes 1 and 2 whereas the other two methods do not (Figure 4b). The first scGBM factor loading can be interpreted qualitatively as saying that moving left in the scGBM factor score plot in Figure 4a corresponds to an increase in gene 1 expression, and moving right corresponds to an increase in gene 2.

**Figure 4.**
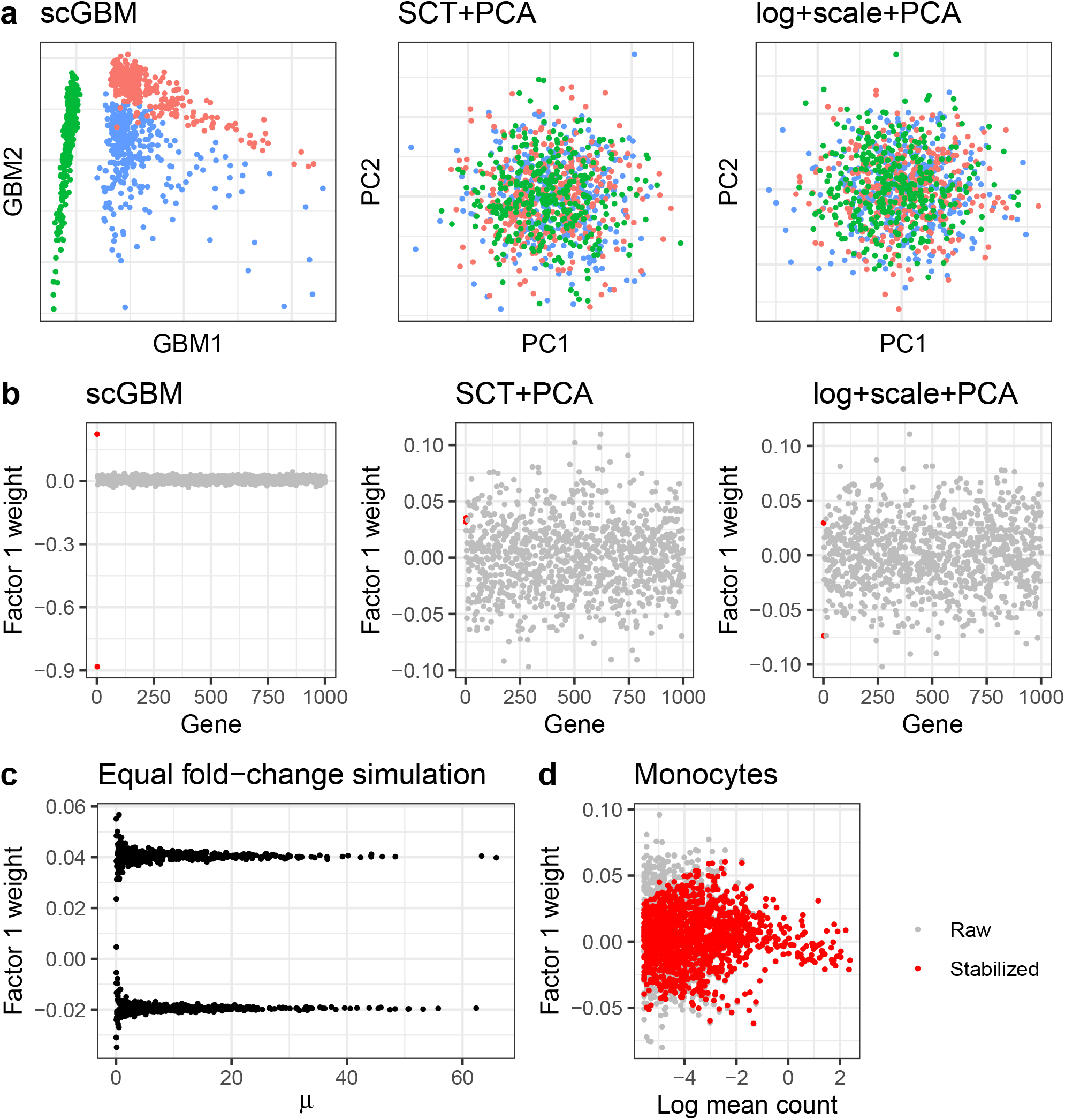
Applying scGBM to the simulations from Figure 1. **a**. Single marker genes simulation: the first two columns of *V*Σ for scGBM compared to embeddings from SCT+PCA and log+scale+PCA. As in Figure 1, cell type A is colored in red, B is colored in green, and C is colored in blue. **b**. The weight of each gene in the first factor. The first two genes (colored red) are the only ones that are truly differentially expressed in the simulation. **c**. Equal log-fold change simulation: Factor 1 weights for scGBM on the simulated data from Figure 1c. **d**. Purified monocytes data: Factor 1 weights for scGBM on the purified monocytes from Figure 1f. The red dots are stabilized weight estimates using adaptive shrinkage based on sampling variability (Section S1.3).

For this simulation, we used the default number of latent factors, *M* = 20, for scGBM. To check for sensitivity to the choice of *M*, we also applied scGBM with *M* = 50 latent factors. The estimated factor scores for *M* = 20 and *M* = 50 agreed well with each other (*r*^2^ = 0.94 for factor 1, *r*^2^ = 0.92 for factor 2). Meanwhile, for GLM-PCA, although it was able to separate the clusters with *M* = 50, we found that the results were highly sensitive to the random initialization used by the algorithm. Occasionally, the first two GLM-PCA scores were very strongly correlated, meaning that the second estimated factor was redundant (Figure S5). In contrast, scGBM is not a randomized algorithm and did not suffer from this issue.

#### Equal log-fold change simulation

Next, consider the equal log-fold change simulation (Figure 1e) where there are two cell types and all genes are differentially expressed with a log_2_ fold change of *±* 1. Unlike SCT+PCA, scGBM does not assign higher weights to genes simply because they have a higher baseline expression (Figure 4c). Indeed, appealingly, the scGBM factor weights are not correlated with the base mean *µ*. GLM-PCA performed similarly to scGBM on this example (Figure S6).

#### Purified monocytes data

As a real data example, we revisit the purified monocytes data (Figure 1f). Here, we find that the GLM-PCA weights are strongly correlated with the log mean count (Figure S6), whereas scGBM does not exhibit this bias. However, we observe that both the scGBM and GLM-PCA weights have a higher variance for genes with low mean, which is undesirable. This is due to the inherent large uncertainty in log(*µ*_*ij*_) when *µ*_*ij*_ is close to zero, leading to unstable estimates of *u*_*im*_ (Figure 4d, raw). To address this, we take advantage of the fact that scGBM is model based, which enables us to quantify uncertainty in the parameter estimates. Using the classical method of inverting the Fisher information matrix (Section 3.5), we approximate the standard error corresponding to each entry of *U* and use the Adaptive Shrinkage (*ash*) method of Stephens (2017) to perform *post hoc* shrinkage towards 0; see Section S1.3 for details. Compared to the usual shrinkage approach based on penalized likelihood, the *ash* method has the advantage of adaptively setting the amount of shrinkage for each parameter, rather than requiring it to be set *a priori*. Since *ash* shrinks genes with larger uncertainties more strongly (Figure 4d, stabilized), this effectively stabilizes the scGBM weights such that low mean genes are not given overly large weights simply due to noise.

#### Semi-simulated hybrid T/B cells

For this example, we used real data to construct a semi-simulated dataset with cells that lie on a “gradient” between two different cell types. Beginning with a dataset of B cells and naive T cells from the *DuoClustering2018* package (Duò et al., 2018), we simulated hybrid cells by combining a fraction of the genes from each cell type. Specifically, we used the following procedure to generate a dataset with *I* = 6168 genes and *J* = 5000 hybrid cells, where each cell is generated as follows:

1. Randomly select one B cell and one naive T cell.
2. Sample *I*^*′*^ *∼* Unif{1, …, *I*} and randomly choose a subset of *I*^*′*^ genes.
3. Construct a simulated cell such that the counts for the *I*^*′*^ randomly chosen genes are equal to those of the memory T cell, and the remaining *I − I*^*′*^ genes counts are equal to those of the naive T cell.

We then tested the ability of each method to capture the biological gradient, which is mathematically represented by the mixture proportion *I*^*′*^*/I* (Figure 5a). Visually, scGBM appears to capture the true latent structure better than SCT+PCA and log+scale+PCA, as both methods appear to be sensitive to the presence of a few outlier cells Figure 5a. The first PC scores from both SCT+PCA and log+scale+PCA were more strongly correlated with the number of UMI counts per cell, suggesting that these methods were less successful at removing the unwanted variability due to sequencing depth (Figure 5b). The spurious groups of cells persist even after uniform manifold approximation (UMAP) is applied to the PCA scores (Figure S7). An additional strength of scGBM is that the low-dimensional embeddings are directly interpretable. Specifically, a unit change along the *x*-axis of the scGBM plot in Figure 5a corresponds to a log fold-change of 1 with respect to the linear combination of genes defined by the first factor loadings. Because scGBM provides a *p*-value for each element *u*_*i*1_, one can visualize the important genes in each factor using a volcano plot (Figure 5c). Theoretical calculations suggest that genes with absolute weight |*u*_*i*1_| greater Than 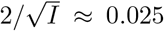should be considered as relevant to the factor (Section S4). For example, *FCRL2* (*u* = *−* 0.035, *p* = 1.28 *×* 10^*−*14^) is known to be preferentially expressed by B cells (Li et al., 2014), whereas *TCEA3* (*u* = 0.028, *p* = 1.53 *×* 10^*−*15^) is reported to be enhanced in naive and memory T cells (Uhlénet al., 2015, Human Protein Atlas, www.proteinatlas.org).

**Figure 5.**
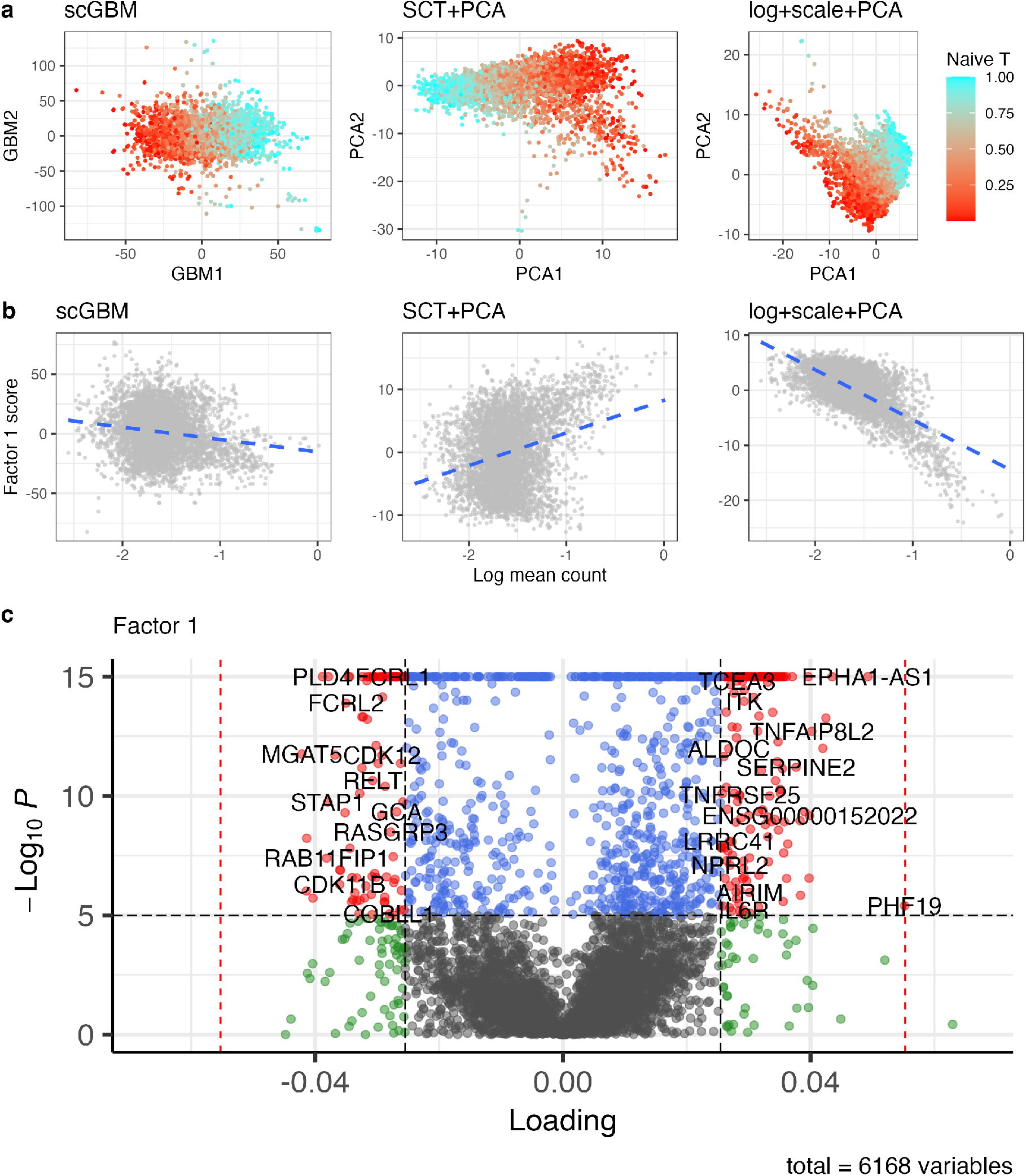
Comparing methods on semi-simulated data where cells are drawn to lie on a gradient between B cells and Naive T cells. **a**. The embeddings from scGBM, SCT+PCA, and log+scale+PCA. Lower values of the mixture proportion (red) indicate that the cell is more like a B cell whereas higher values (light blue) indicate that the cell is more like a Naive T cell. **b**. Factor 1 score versus the log of the mean UMI count per cell. **c**. A “volcano” plot of the genes driving the first scGBM factor. Specifically, the loading weight for each gene is plotted against its*−* log_10_ p-value. This plot was made using the *EnhancedVolcano* R package (Blighe et al., 2023).

### 4.4 Uncertainty quantification and the cluster cohesion index

Clustering typically occurs downstream of dimensionality reduction, as a *post hoc* analysis step (Kiselev et al., 2019). Uncertainty in the estimated low-dimensional representation is likely to influence clustering results and other downstream analyses, but the effect of this uncertainty has not been thoroughly investigated in previous work. For example, one might wonder whether different clusters would be identified if the same set of cells was re-sequenced and the same analysis was performed on the resulting new count matrix.

To illustrate, we simulated a count matrix consisting of random Poisson noise, *Y*_*ij*_ *∼* Poisson(1) i.i.d. for *i* = 1, …, *I* and *j* = 1, …, *J* with *I* = 1000 and *J* = 5000, and we used scGBM to obtain estimated score 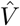. Then we applied the Louvain algorithm (Blondel et al., 2008) to 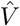 to identify clusters of cells, since this is the default clustering algorithm used in Seurat (Satija et al., 2015) (FindClusters() with default resolution equal to 0.8). The Louvain algorithm identified 9 clusters, even though the cells were simulated to be homogeneous. The fact that standard single-cell clustering algorithms produce too many clusters on null data such as this has also been noted in other studies (Morelli et al., 2021; Grabski et al., 2022).

With scGBM, we can use the uncertainty in 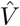 to quantify the uncertainty in these clusters. First, a simple visualization of the uncertainty in 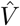 can be obtained by drawing an ellipse around each point, with the dimensions of the ellipse equal to the estimated standard errors; see Figure 6a (left). The considerable overlap among these ellipses suggests that there are no clearly separated clusters. Meanwhile, when there are true clusters, the ellipses for cells from different clusters tend not to overlap as much. For example, on the 10X immune cell data, there is relatively little overlap between the uncertainty ellipses for the distinct cell populations (T cells, B cells, monocytes); see Figure 6a (right). However, the qualitative visualizations in Figure 6a may be difficult to interpret or use, especially when there are many clusters.

**Figure 6.**
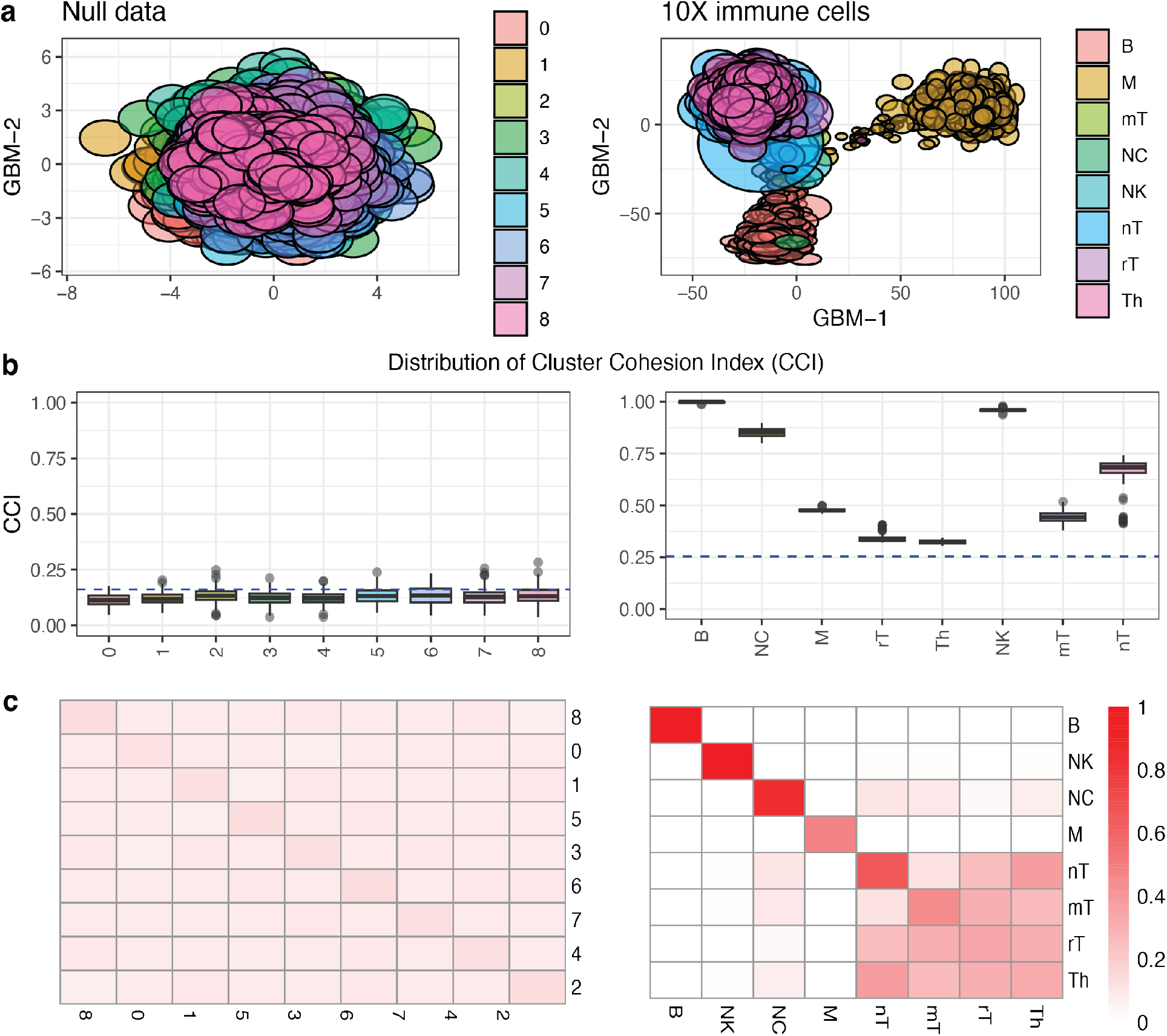
Cluster cohesion results for two datasets: random Poisson noise and the 10X immune cells. In the 10X immune data, B = “B cells”, M =“Monocytes”, mT = “Memory T”, NC = “Naive cytotoxic”, NK = “Natural Killer”, nT = “Naive T”, rT = “Regulatory T”, and Th = “T helper”. **a**. The first two columns of 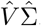. The uncertainty in each point is visualized by drawing an ellipse with axis lengths equal to the estimated standard errors. **b**. The distribution of the CCI fractions *f*_*k,k*_; see Equation S1.16. **c**. Heatmaps of inter-CCIs for the random Poisson noise data and the 10X immune cells dataset. The inter-CCIs enable one to visualize the uncertainty in the distinctions between different clusters.

#### Cluster cohesion index (CCI)

To provide a quantitative measure of the uncertainty in each cluster, we introduce the *cluster cohesion index* (CCI). Given estimates 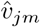 and corresponding standard errors 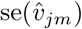, we randomly generate perturbed Values 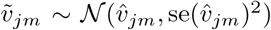 independently, perform clustering on the rows of 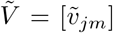, and compute the fraction of pairs of cells that are still in the same cluster. Then, for each original cluster, the CCI is defined as the mean of this fraction over many repetitions; see Section S1.4 for details. The CCI takes a value between 0 and 1, with 1 indicating that all pairs of cells are still in the same cluster after resampling.

Figure 6b shows the CCIs for the random Poisson noise dataset and the 10X immune cell dataset from Figure 6a. As expected, the CCIs for the random noise dataset are all low, reflecting the fact that any identified clusters are artifacts of sampling variability. Meanwhile, on the 10X immune cell data, the CCIs for the true cell types are all higher. We can define a significance threshold for CCIs by computing the values that would have been generated under the null model of *V* = 0; see Section S1.4 for details. As one would hope, all of the random Poisson noise CCIs fall below the significance threshold, while all of the 10X immune cell CCIs are above the threshold; see dashed blue lines in Figure 6b.

### Inter-cluster cohesion index

To quantify the overlap between clusters in a way that accounts for model-based uncertainty, we define the *inter-cluster cohesion index* (inter-CCI) as follows: out of all pairs of points *j* and *j*^*′*^ that were originally in clusters *k* and *k*^*′*^, respectively, the inter-CCI is the mean of the fraction that are in the same cluster after resampling; see Section S1.4 for details. Note that the inter-CCI between a cluster and itself coincides with the CCI of that cluster.

The inter-CCIs can be visualized using a heatmap (Figure 6c). For the random Poisson noise data, all of the inter-CCIs are of similar magnitude to the CCIs themselves, indicating low confidence in all of the clusters, as expected. On the 10X immune cell data, the inter-CCIs for the T-cell subpopulations (nT, mT, rT, Th) are also relatively high, indicating lower confidence in the distinction between these groups. Meanwhile, the remaining subpopulations (B, NK, NC, and M) have high CCIs and low inter-CCIs, indicating high confidence in these clusters. The inter-CCIs allow one to visualize and quantify cluster relationships that would be difficult to discern using scatterplots alone.

### Application to breast cancer data

To further demonstrate the utility of inter-CCIs, we consider a dataset of 100,064 cells from the tumor microenvironment of 26 breast cancer patients (Wu et al., 2021); see Figure 7a. Since Wu et al. (2021) used the standard Seurat pipeline (Satija et al., 2015) to cluster cells and used Garnett (Pliner et al., 2019) for annotation, we computed the inter-CCIs for this Seurat-based clustering; see heatmap in Figure 7b. The blocks along the diagonal indicate groups of cell types with substantial overlap, accounting for model-based uncertainty. These blocks appear to correspond to biologically relevant classes of cells such as lymphoid (T cells, B cells, NK cells), myleoid (monocytes and DCs), and endothelial cells. The inter-CCI heatmap enables us to see the relationships among all of the 29 minor cell types, which would be difficult or impossible to see with a scatterplot.

**Figure 7.**
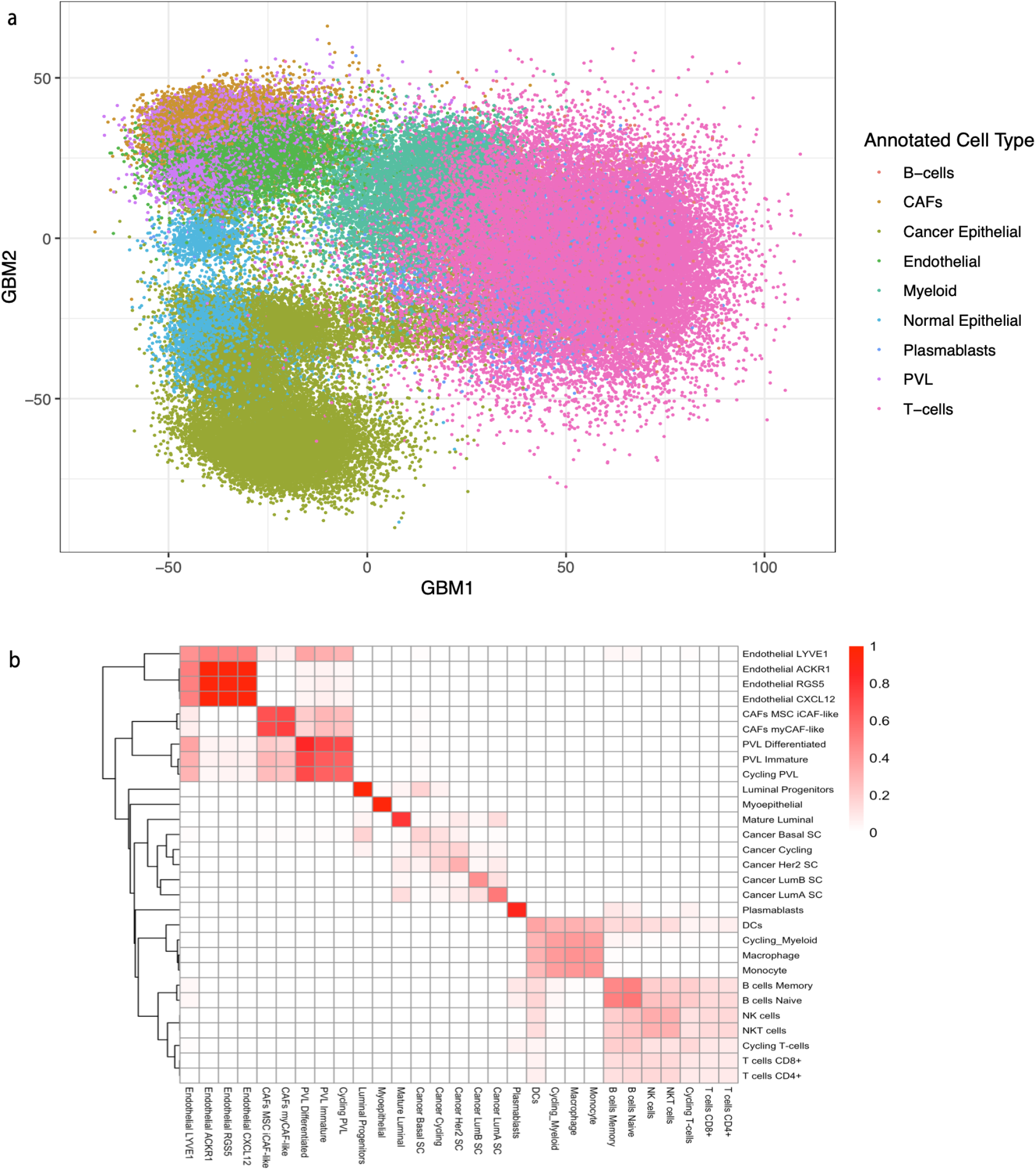
**a**. We applied scGBM to the Wu et al. (2021) dataset consisting of 100,064 breast cancer cells and plotted the first two scGBM scores (columns of 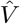) colored by major cell type. **b**. The heatmap of inter-cluster cohesion indices (inter-CCIs) for the minor cell types. The inter-CCIs clearly reveal relationships between the clusters that are not visible in the scatterplot, and even enable one to analyze the numerous minor cell types, which would be very difficult to visualize in a 2-d scatterplot

## 5 Discussion

Currently, standard practice is to visualize single-cell data by applying UMAP or t-SNE (McInnes et al., 2018; Van der Maaten and Hinton, 2008) to the PCA scores. Although UMAP could instead be applied to the scores produced by scGBM, our results from Figures 4 and 5 show that the raw scGBM scores *V Σ* are already sufficient for visualizing the dominant sources of biological variability. An additional advantage of scGBM is that the visualizations have a direct biological interpretation by looking at the genes included in the corresponding factors. Since UMAP embeddings been shown to induce undesirable distortions (Chari and Pachter, 2023), we believe that scGBM can provide a useful alternative for users who prefer a non-linear yet still model-based embedding.

In future work, there are several interesting directions for improving upon our current approach. Although our scGBM algorithm is faster than GLM-PCA, it is still much slower than PCA — especially because commonly used transformations preserve sparsity, which allows for the use of more efficient SVD algorithms (Baglama and Reichel, 2005). This limitation could potentially be addressed by modifying our IRSVD algorithm to avoid applying the SVD to a dense matrix. A second limitation of scGBM is that, unlike PCA, the latent factors depend on the choice of *M*. Although there are heuristics for choosing the number of latent factors *M*, developing principled techniques for choosing *M* remains an area for future work. In this paper, we mainly considered quantifying uncertainty in the scores *V*, but it would be also be interesting to develop additional approaches that use the uncertainty in the gene loadings *U*. For example, one could propagate uncertainty into gene set enrichment analysis (Subramanian et al., 2005). A final area for future research would be to induce sparsity in the weights *U*, to improve biological interpretability. Sparsity in *U* would mean that only a few genes have non-zero weights, making it easier to connect the factors to known biology, for instance, in terms of gene sets and pathways. To this end, it would be interesting to explore whether ideas from sparse PCA (Zou et al., 2006) can be extended to bilinear models.

## Acknowledgments

The authors thank Will Townes for helpful comments and suggestions. PBN is supported by the National Institutes of Health grant T32CA009337. JWM is supported by the National Institutes of Health grant 5R01CA240299.

## Supplementary Material for

### S1 Extended Methods

#### S1.1 Generalized bilinear model

The scGBM method employs the Poisson bilinear model in Equation 3.1 for the matrix *Y* ∈ ℝ^*I×J*^ of UMI counts (*I* genes and *J* cells). If we define *μ*:= [*µ*_*ij*_] ∈ ℝ^*I×J*^ then Equation 3.1 can be rewritten in matrix form as

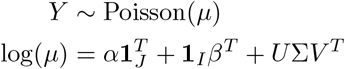

where Poisson and log are applied entry-wise, **1**_*K*_ = (1, …, 1)^*T*^ ∈ ℝ ^*K*^ is a vector of ones, and Σ = diag(*σ*_1_, …, *σ*_*M*_) ∈ ℝ ^*M ×M*^. Theorem 5.1 of Miller and Carter (2020) shows that this model is identifiable under the following constraints:

- *U*^*T*^ *U* = *V* ^*T*^ *V* = *I*_*M*_ (orthonormality),
- *σ*_1_ > … > *σ*_*M*_ *>* 0,
- the first non-zero entry in every column of *U* is positive,
- 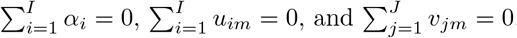

Many of the results presented in this section can easily be extended to cases with more complicated experimental designs (for example, adding row and column covariates), different link functions, and different outcome distributions. See Miller and Carter (2020) for more general GBM formulations.

#### S1.2 Iteratively reweighted singular value decomposition (IRSVD)

##### Weighted low-rank approximation

Given a matrix *Z ∈ ℝ*^*I×J*^, the weighted low-rank problem is to find a rank *M* matrix *X∈ ℝ*^*I×J*^ that minimizes the weighted squared error between *X* and *Z*:

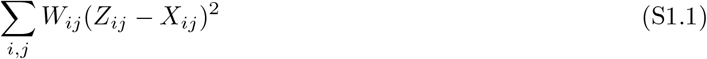

where *W ∈ ℝ* ^*I×J*^ is a matrix of known non-negative weights. When *W*_*ij*_ = 1 for all *i* and *j*, a solution can be found via the truncated singular value decomposition (SVD) of *Z* (Eckart and Young, 1936). Unfortunately, the general case cannot be reduced to an eigenvector problem unless rank(*W*) = 1. When the weights have been scaled to be in [0, 1], Srebro and Jaakkola (2003) present the following iterative algorithm to find *X*:

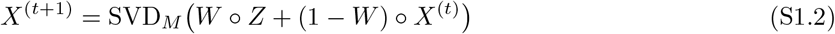

where ° is the Hadamard product and SVD_*M*_ (*X*) denotes the rank *M* truncated SVD of *X*. Tuzhilina and Hastie (2021) note that Equation S1.2 can be seen as a projected gradient descent step: since the gradient is

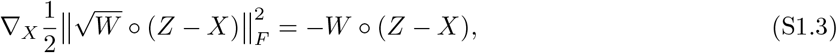

a gradient step would be

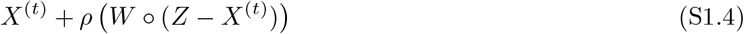

which coincides with the argument of SVD_*M*_ in Equation S1.2 when the step size is *ρ* = 1. The projection back onto the set of rank *M* matrices is given by SVD_*M*_. Further, Tuzhilina and Hastie (2021) show that the convergence rate can be improved by using the acceleration method of Nesterov (1983):

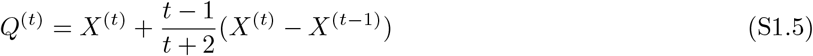

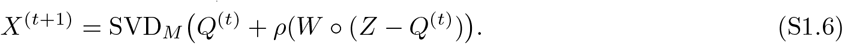

##### Approximating the Poisson bilinear model log-likelihood

We show that maximum likelihood estimation for *X* in the Poisson bilinear model in Equation 3.1 can be locally approximated by a weighted low-rank problem. Denoting *X*_*ij*_ = (*U*Σ*V* ^*T*^)_*ij*_, the log-likelihood of *X* is

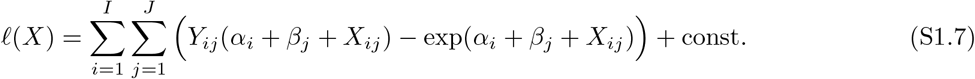

For now, we hold *α*_*i*_ and *β*_*j*_ at arbitrary fixed values. Differentiating with respect to *X*_*ij*_, we have

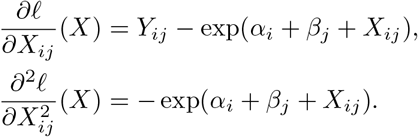

Suppose 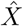 is our current estimate of *X*, and denote 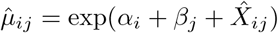. Then a second-order Taylor approximation at 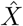yields

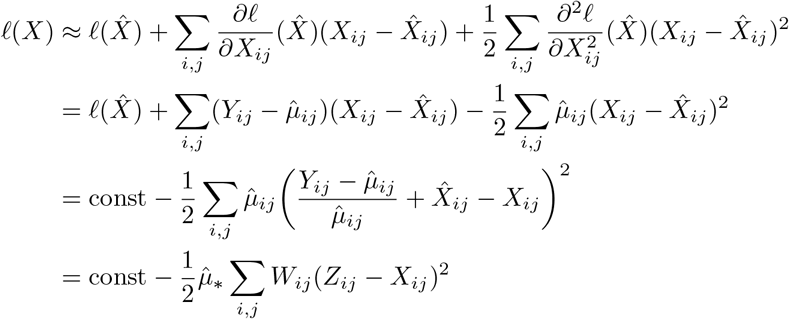

where

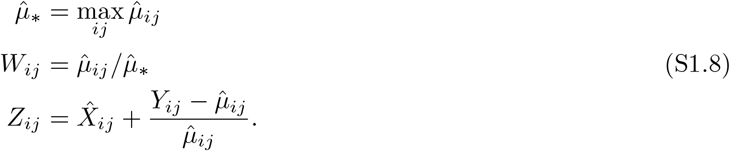

Under this local approximation,

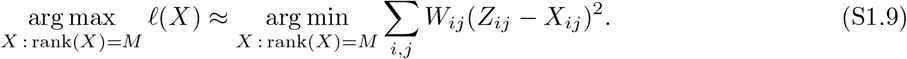

Thus, maximizing the likelihood of *X* in the Poisson bilinear model can be locally approximated by a weighted low-rank problem of the form in Equation S1.1, which can be solved using the Srebro and Jaakkola (2003) iteration in Equation S1.2.

##### Iteratively reweighting based on the local approximations

Unlike the standard weighted low-rank problem, however, the *W* and *Z* in Equation S1.9 depend on the current parameter estimates rather than being fixed. This suggests using an iterative algorithm in which the local approximation is sequentially updated based on the current estimates of 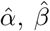, and 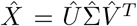. Specifically, we propose alternating between the following two steps.

###### 1. Update intercepts

Holding 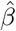 and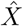 fixed, for each *i*, the model reduces to a standard GLM with intercept *α*_*i*_. In the Poisson case, there is a closed-form solution for maximizing with respect to *α*_*i*_, which yields the update in Equation 3.5. Likewise, if one wishes to estimate *β*_*j*_, this can be done in the same way by maximizing with respect to *β*_*j*_, yielding Equation 3.6.

###### 2. Update latent factors

Given estimates 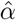 and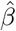, along with our current estimate 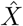, define *W*_*ij*_ and *Z*_*ij*_ by Equation S1.8. To obtain a new 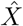 that is closer to solving the local approximation in Equation S1.9, apply a generalized form of the Srebro and Jaakkola (2003) iteration by combining Equations S1.2 and S1.4, which yields the update in Equation 3.7. This can also be modified to use Nesterov acceleration as in Equation S1.5.

##### Initialization

Using a good choice of initialization is important for fast and reliable convergence. If we were to initialize 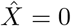, then after the first update to the intercepts, the weights would be

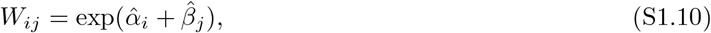

which makes *W* a rank 1 matrix. When rank(*W*) = 1, it turns out that the weighted low-rank problem can be solved in one step by reducing to a standard SVD. Thus, it is preferable to solve this initial weighted low-rank problem exactly rather than just performing one iteration of Equation 3.7. The exact solution is derived as follows.

###### Proposition S1.1

*If* 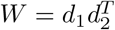 *Where* 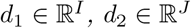 *for all i, j, then*

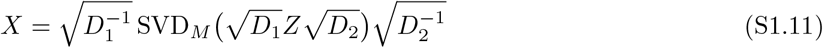

*minimizes Equation* S1.1 *subject to* rank(*X*) *≤ M, where D*_1_ = diag(*d*_1_) *and D*_2_ = diag(*d*_2_).

*Proof*. The proof is due to Razenshteyn et al. (2016). Since *W*_*ij*_ = *d*_1*i*_*d*_2*j*_, we have

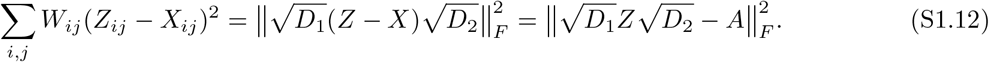

where 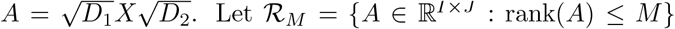 Let ℛ_*M*_ = {*A* ∈ ℝ^*I×J*^ : rank(*A*) *≤ M*}. By the Eckart–Young theorem, 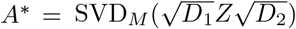 minimizes Equation S1.12 over *A ∈ ℛ*_*M*_. Since *W*_*ij*_ *>* 0 implies *d*_1*i*_ *> 0* and *d*_2*j*_ *>* 0, it follows that 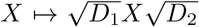is a bijection from *ℛ*_*M*_ to itself. Thus, 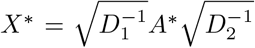 Equation S1.12 over *X ∈ ℛ*_*M*_.

Applying Proposition S1.1 suggests using an initial estimate of

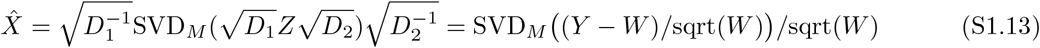

where sqrt and */* denote entry-wise square root and entry-wise division, respectively. In practice, it is necessary to clip extreme values of 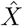 before proceeding, since extreme values can lead to instability in the SVD. Thus, we hard threshold using *x →* max(min(*x, c*), *−c*) to ensure all values of 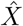 are in [*−c, c*]. By default, we use *c* = 8. This clipping procedure is similar to the one used by scTransform (Hafemeister and Satija, 2019). Combining this clipping procedure with Equation S1.13, we arrive at our proposed initialization procedure in Section 3.3.

Interestingly, note that since 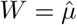, the entries of (*Y − W*)*/*sqrt(*W*) coincide with the Pearson residuals under a Poisson model. Thus, the derivation of Equation S1.13 provides some theoretical justification for using PCA on the Pearson residuals, as done by scTransform (Hafemeister and Satija, 2019), by thinking of it as an approximation to GBM parameter estimation. However, the assumption that *W* is rank 1 (or close to it) is unlikely in cases where there is a large amount of latent structure. Thus, in many cases of interest, this will only be a very rough approximation.

#### S1.3 Adaptive shrinkage for stabilization

Miller and Carter (2020) and Townes (2019) use *ℓ*_2_ penalties on the factors *U* and *V* to stabilize the estimates via shrinkage towards 0. However, this requires a tuning parameter to be chosen in advance. Instead, for scGBM, we propose to use the following *post hoc* adaptive shrinkage method for the weights *U*, based on the *ash* method of Stephens (2017).

Let *Û* ∈ ℝ^*I×M*^ be the estimate of the *U* matrix, and let *s*_*im*_ := se(*Û* _*im*_) be the estimated standard error of entry *Û*_*im*_. Consider the following spike-and-slab model for *U*, introduced by Stephens (2017):

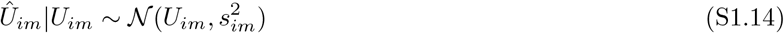

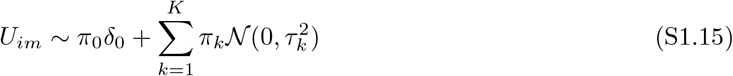

for each *i* and *m* independently, where *δ* _0_ denotes the point mass at zero. The component variances 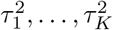 are pre-specified and the mixture weights *π*_0_, …, *π*_*K*_ are estimated using an empirical Bayes procedure. The idea is that the true parameter value *U*_*im*_ is drawn from a spike-and-slab distribution and the empirical estimate *Û*_*im*_ is drawn from a normal distribution with mean *U*_*im*_ and standard deviation equal to the estimated standard error of *Û* _*im*_. The posterior on *U*_*im*_ given *Û*_*im*_ is then used for uncertainty quantification. The method is called *adaptive shrinkage* and is implemented in the R package *ashr* (Stephens et al., 2022). In Figure 4d, the stabilized estimates are defined as the posterior mean of *U*_*im*_ given *Û*_*im*_ under the *ash* model.

#### S1.4 Cluster cohesion index

Given estimates 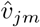 and corresponding standard errors 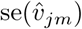, we aim to quantitatively analyze the stability of a given clustering of cells. Let *c*_1_, …, *c*_*J*_ 1, …, *K* be assignments of the *J* cells to *K* clusters. To compute the *cluster cohesion indices* (CCIs), we repeat the following steps *n* times (by default, *n* = 100):

1. Draw 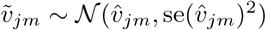for *j* = 1, …, *J, m* = 1, …, *M*.
2. Apply a clustering algorithm to the rows of 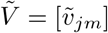to obtain new cluster assignments 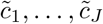.
3. For each pair of clusters *k, k*^*′*^ *∈ {*1, …, *K}*, compute the fraction

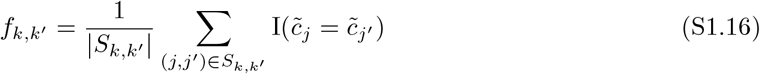

where *S*_*k,k*_*′* is the set of pairs *j, j*^*′*^ ∈ {1, …, *J*} such that *j* ≠ *j*^*′*^, *c*_*j*_ = *k*, and *c*_*j*_*′* = *k*^*′*^.

In Equation S1.16, I(*·*) denotes the indicator function. Then, for each cluster *k*, the CCI is defined as the mean (across the *n* repetitions) of the fraction *f*_*k,k*_. Likewise, the inter-cluster cohesion index (inter-CCI) for clusters *k* and *k*^*′*^ is defined as the mean of *f*_*k,k*_*′*. That is, the inter-CCI is the mean of the fraction of all pairs of points *j* and *j*^*′*^ that are in the same cluster after resampling, out of all pairs *j* and *j*^*′*^ that were originally in clusters *k* and *k*^*′*^, respectively.

The interpretation is that a low CCI indicates that this cluster could be an artifact of sampling variability. Likewise, a high inter-CCI for clusters *k* and *k*^*′*^ indicates that the separation of these two clusters may be an artifact of sampling variability. The clustering algorithm in step 2 can be specified by the user, and does not have to be the same as the algorithm that was used to create the original assignments *c*_1_, …, *c*_*J*_.

To characterize the CCI under the null of no latent variability we replace 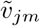 in step 1 with 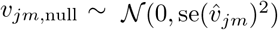 for *j* = 1, …, *J, m* = 1, …, *M*. Then we perform steps 1-3 above for *n* = 100 repetitions, and compute the 95th percentile of the resulting *f*_*k,k*_ values to get the dashed blue lines in Figure 6b.

#### S1.5 Controlling for known batches

When there are known batches, the model can be adjusted to have a gene-level intercept for each batch:

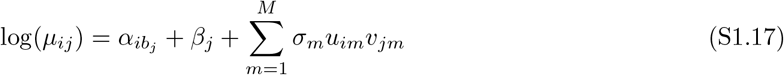

where *b*_*j*_ *∈ {* 1, …, *B}* is the index of the batch that cell *j* belongs to, and *α*_*ib*_ is the intercept for gene *i* and batch *b*. Estimation in this model is the same as before, except Equation 3.5 is replaced with

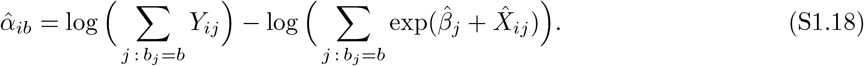

### S2 Data and software availability

We briefly describe the single-cell datasets used in the paper. Unless otherwise specified, all single-cell data was processed using Seurat (Satija et al., 2015). When it is stated that variable genes were selected, this means that the function `FindVariableFeatures()` was used to select the genes as input to scGBM.

- The purified monocyte data was downloaded from the 10X genomics website. The count matrix is available in the folder “Gene / cell matrix (filtered).”
- The 10X immune cell data (Zheng et al., 2017), including the Naive T cells and memory T cells used in Figure 5, were downloaded from the *DuoClustering2018* R package (Duò et al., 2018).
- The COVID-19 Atlas (Wilk et al., 2020) was downloaded as a Seurat object from www.covid19cellatlas.org.
- The 10X mouse brain data was downloaded from the Bioconductor package *TENxBrain* (Lun and Morgan, 2020).
- The Wu et al. (2021) data was downloaded from the Gene Expression Omnibus (GEO) database (GSE176078).

scGBM is available for download as an R package at https://github.com/phillipnicol/scGBM. The repository also includes scripts for replicating the figures based on simulated data.

### S3 Technical details of simulations

#### Single marker genes simulation

We generated *J* = 1000 cells such that 333 are from cell type A, 333 are from cell type B, and 334 are from cell type C. For gene 1, counts are independently generated as Poisson(10) for cell type A and Poisson(1) for all other cells. For gene 2, counts are independently generated as Poisson(50) for cell type B and Poisson(1) for all other cells. For genes 3, 4, …, 1000 (the noise genes), all counts are independently generated as Poisson(1). Figure 1a shows boxplots of the simulated counts.

#### Projection method simulation

To test the accuracy of the projection method on a simulation with known ground truth, we simulated latent factors and data as follows with *I* = 1000, *J* = 10000, and *M* = 10. Following the procedure described in Section S2 of Miller and Carter (2020), the ground truth matrices *U* and *V* were drawn uniformly from the rank *M* Stiefel manifolds over ℝ^*I*^ and ℝ^*J*^, respectively. The intercepts *α*_*i*_ and *β*_*j*_ were drawn indendpently from 𝒩 (0, 1). The diagonal elements of Σ were uniformly spaced between 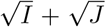 and 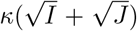, where *κ* is a parameter that controls the amount of latent variability. Miller and Carter (2020) use *κ* = 2, which we call low latent variability (LLV). We also use *κ* = 5 and call this high latent variability (HLV).

We then compared scGBM-full to scGBM-proj (projection method) using various subsample sizes in the projection method, using *M* = 10 for both methods. Specifically, we considered subsample sizes ranging between 100 and 5000 (1%-50%). For each subsample size, we ran both methods on 100 trial experiments, where each trial was performed by simulating a full dataset with *I* = 1000 and *J* = 10000 and choosing a random subsample of the desired size. We assessed the results using two metrics. First, we considered the relative mean squared error between the estimated and ground truth *V* :

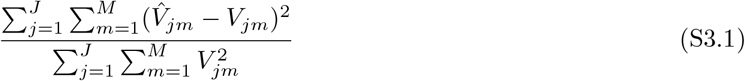

as defined on page S12 of Miller and Carter (2020). We also considered the absolute value of the correlation between columns of true and estimated *V*, as used above in Figure 3. The results are shown in Figure S3.

### S4 Additional calculations

#### Determining genes that are relevant to a given factor

Under the null model, the first estimated factor *Û*_1_ ∈ ℝ^*I*^ is uniformly distributed on the surface of the ball *S*^*I−*1^ := {*x ∈ ℝ*^*I*^ : ||*x*||_2_ = 1}. A heuristic cutoff is to choose genes such that their weight is greater than what would be expected under this null model.

In this case *Û*_1_ has same distribution as a multivariate standard normal random variable *Z∼ N* (0, I) that has been projected onto *S*^*I−*1^:

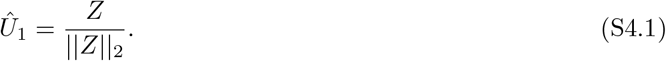

By the law of large numbers and the continuous mapping theorem,

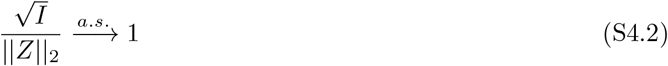

so the distribution of *Û*_1_ can be approximated by the distribution of (suitably scaled) folded normal distributions

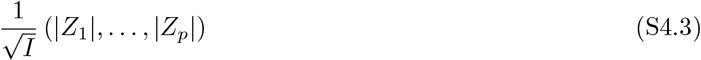

Applying standard results shows that and

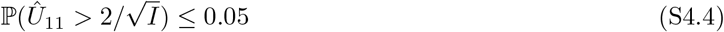

and

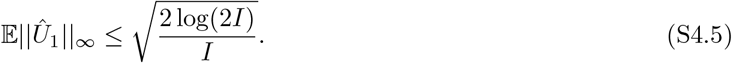

This justifies the use of 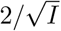 or the more stringent 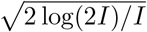 cutoff to consider a gene as relevant to a factor.

### S5 Supplementary figures and tables

**Table S1:** **Sensitivity of GLM-PCA (Avagrad) to learning rate**. Using GLM-PCA (Townes and Street, 2020) with optimizer=“Avagrad” on simulated data with *I* = *J* = 1000 and *M* = 10, the GLM-PCA objective function (Deviance) after 100 iterations is reported for several choices of the learning rate, that is, the step size for gradient descent. If the learning rate is too large, then the algorithm diverges, indicated by a value of *∞*. If the learning rate is too low, then the rate of convergence is very slow. In practice, it may be difficult to find a learning rate that works well for a particular dataset, and thus, the runtime of GLM-PCA can be significantly longer than what is reported in Table 1.

| Learning rate | Deviance (after 100 iterations) |
| --- | --- |
| 0.1 (Default) | ∞ |
| 0.03 | ∞ |
| 0.02 | $9.61 \times 10^5$ |
| 0.01 | $9.51 \times 10^5$ |
| 0.001 | $1.18 \times 10^6$ |

**Figure S1:**
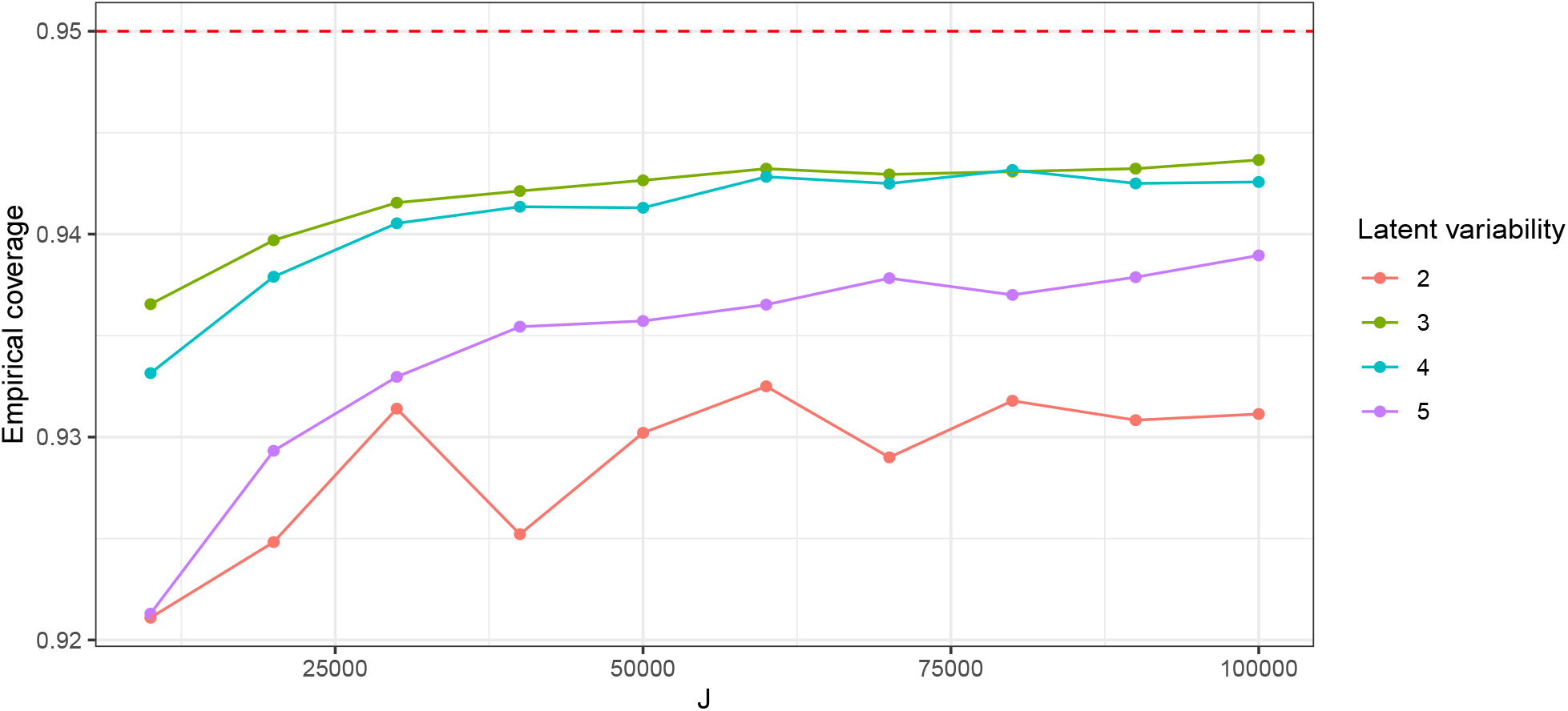
Empirical coverage of 95% confidence intervals across different simulation settings for *J* and latent variability *κ* = Σ_1_*/Σ*_*M*_. Confidence intervals were formed using a normal approximation: 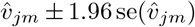. The empirical coverage is the fraction of times the confidence interval contained the true *v*_*jm*_. Points shown are the median across 20 replicates

**Figure S2:**
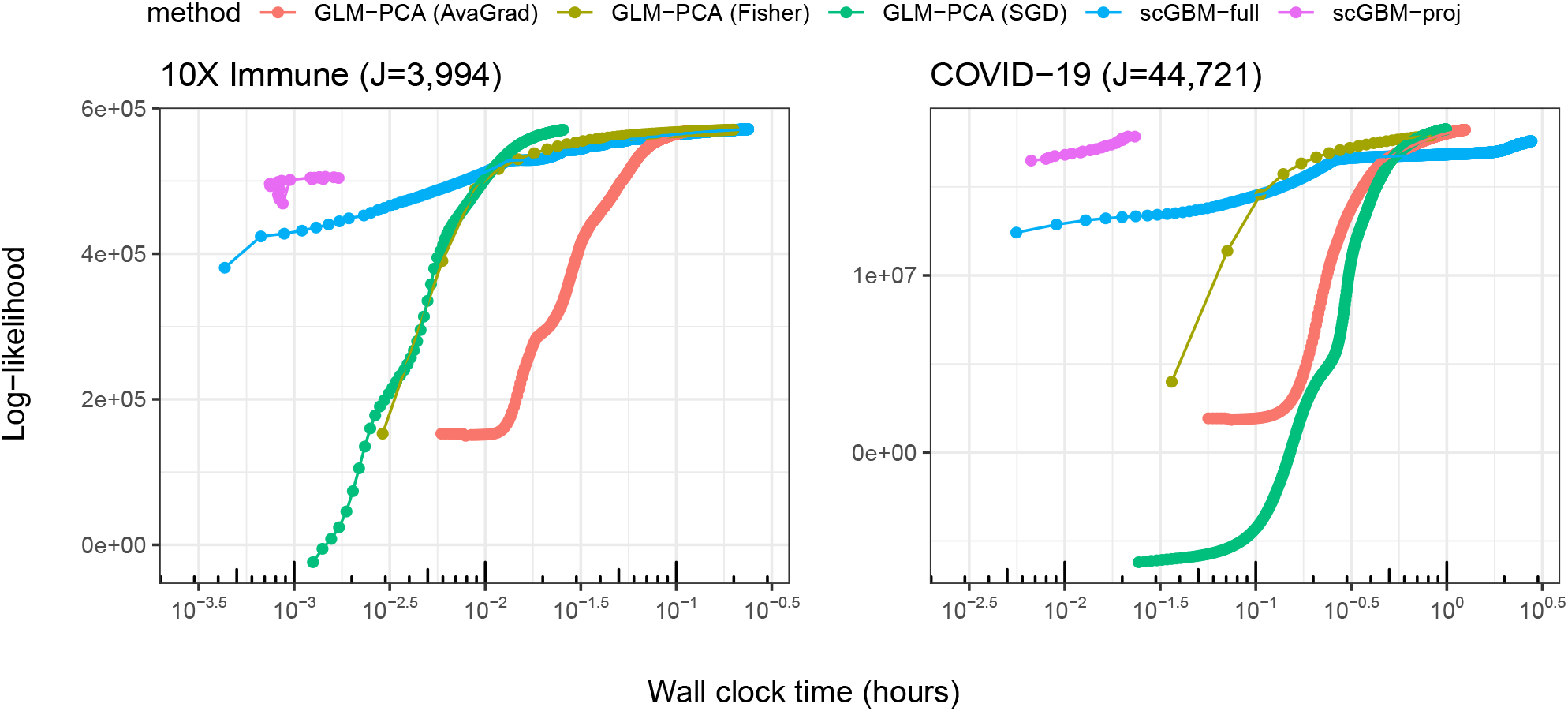
The same analysis as in Figure 2c,d for two real datasets from Table 1.

**Figure S3:**
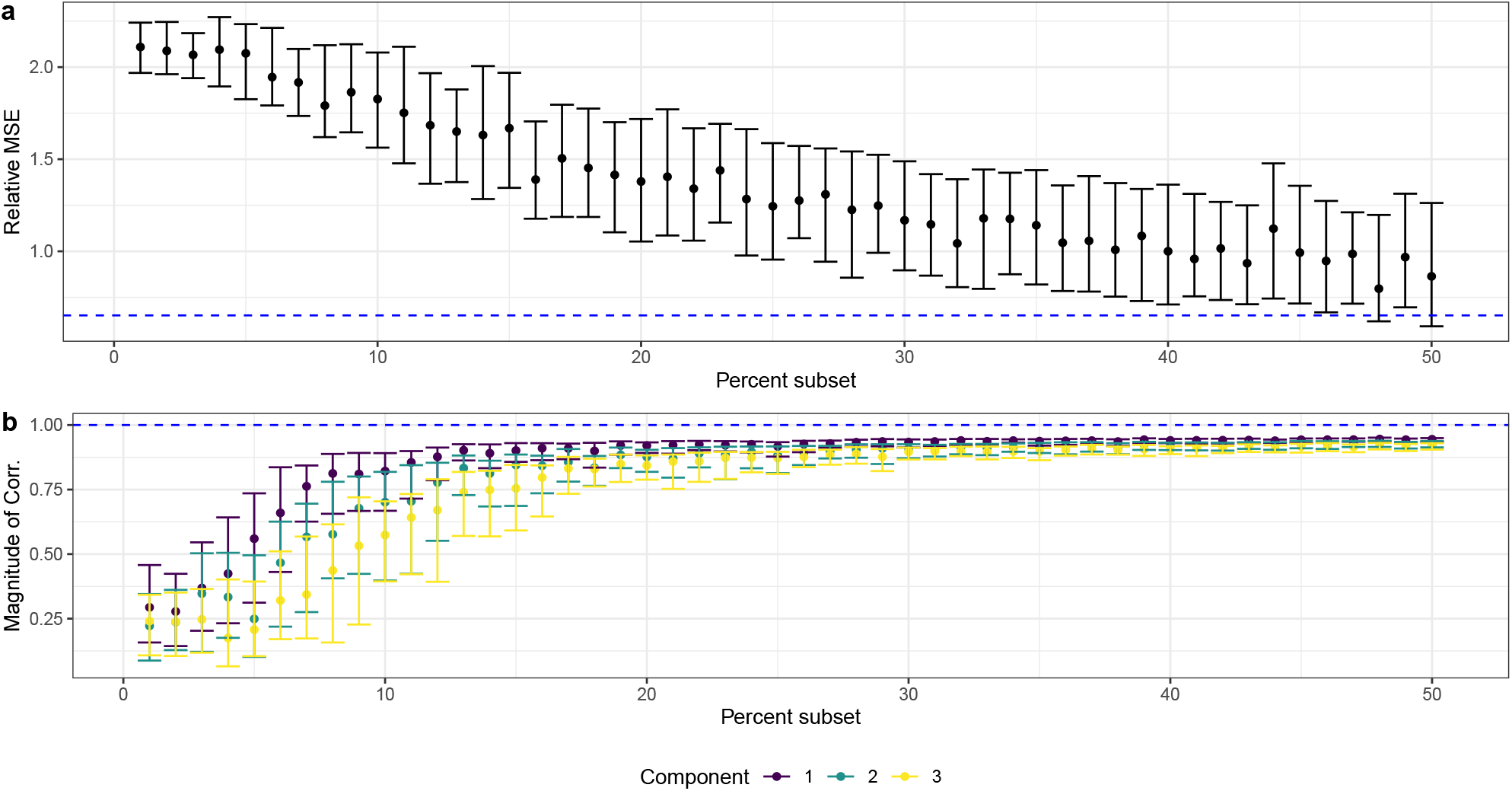
Testing the accuracy of the projection method using simulated data with *I* = 1000, *J* = 10000, and *M* = 10. **a**. Relative MSE between ground truth and scGBM-proj estimate of *V* as a function of subsample size used. The points are the median across 100 trials and the error bars represent the interquartile range. The dashed blue line is the median of the relative MSE for scGBM-full. **b**. Absolute value of the correlation between ground truth and scGBM-proj estimate of *V* as a function of subsample size. This is a simulation-based replication of the real data analysis in Figure

**Figure S4:**
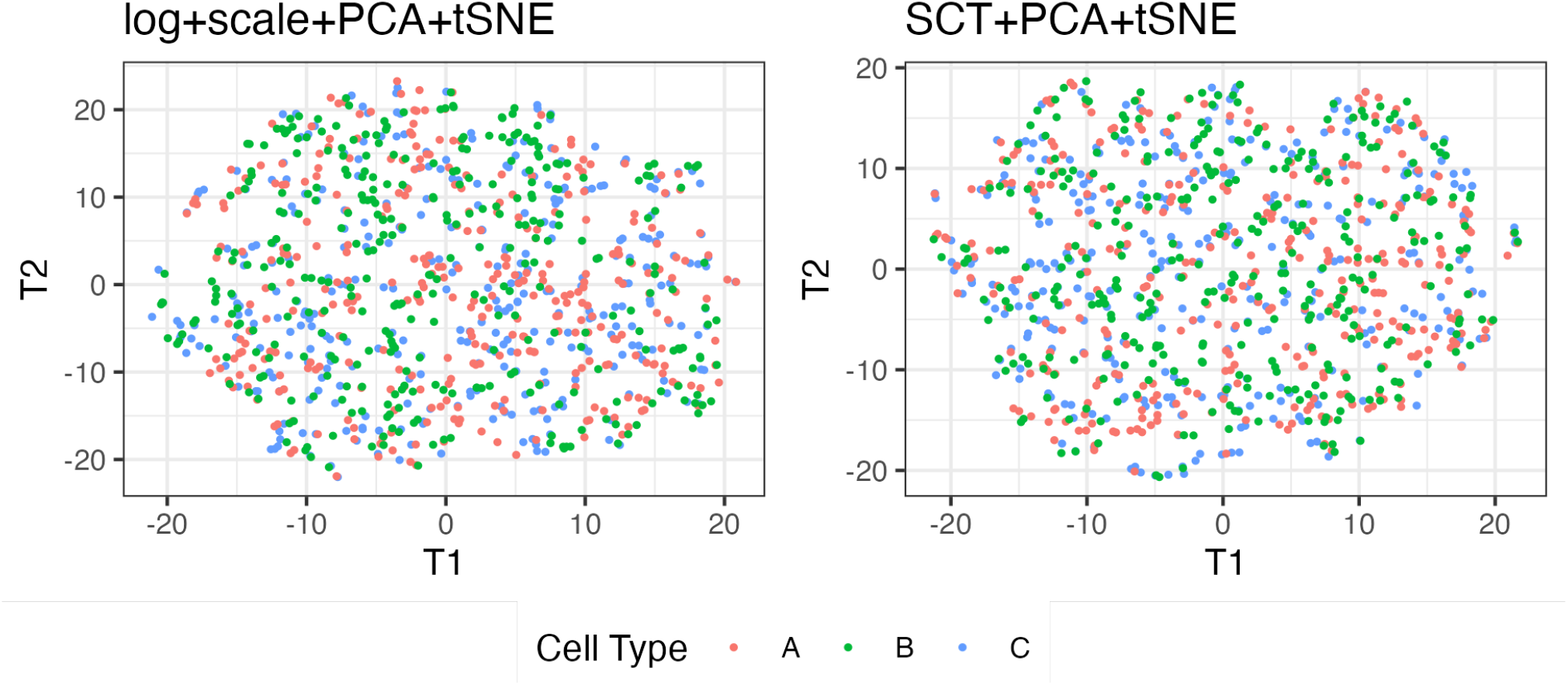
Results from applying t-SNE (Van der Maaten and Hinton, 2008) to the output of SCT+PCA and log+scale+PCA on the single-marker gene simulation; compare with Figures 1 and 4.

**Figure S5:**
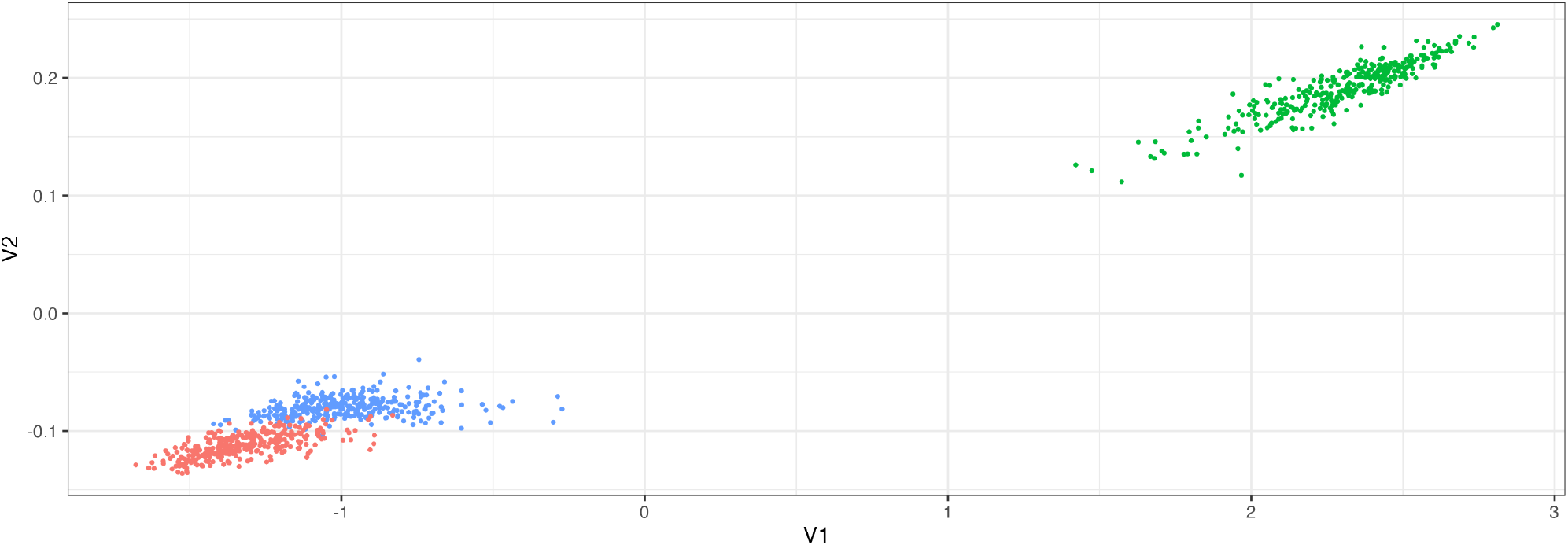
Results from applying GLM-PCA (AvaGrad) to the single-marker gene simulation from Figures 1 and 4. The scores for factors 1 and 2 lie close to a line, which suggests that the first two factors are essentially the same.

**Figure S6:**
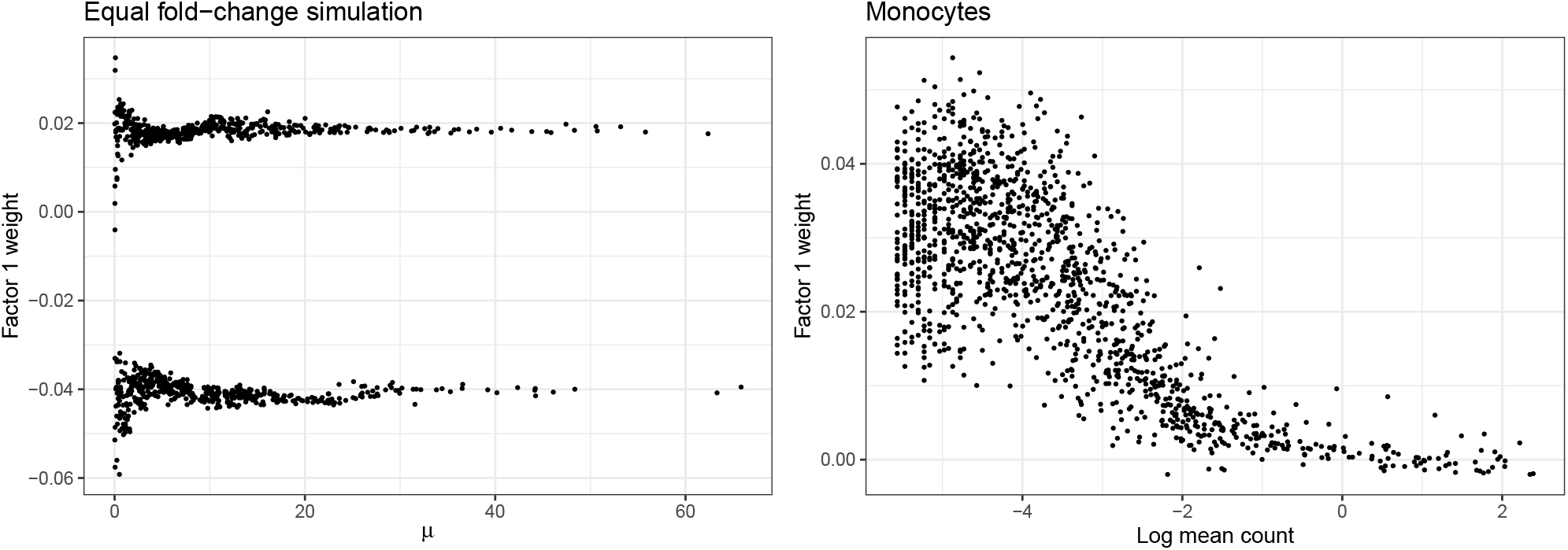
Results from applying GLM-PCA (AvaGrad) to the equal log-fold change simulation and the purified monocytes dataset; compare with Figure 1e,f and Figure 4c,d.

**Figure S7:**
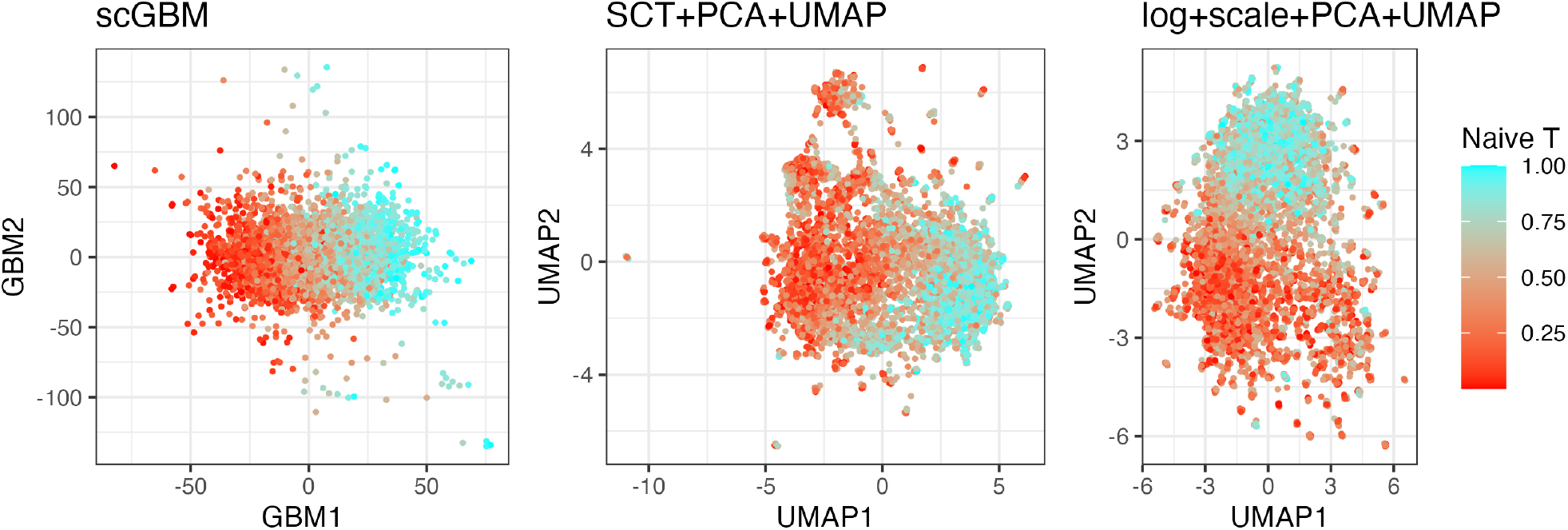
Results from applying UMAP (McInnes et al., 2018) to the output of SCT+PCA and log+scale+PCA on the semi-simulated hybrid T/B cell data; compare with Figure 5.

## Footnotes

1 https://github.com/phillipnicol/scGBM

## Notes

https://github.com/phillipnicol/scGBM

### Competing Interest Statement

The authors have declared no competing interest.

### Summary of Updates

Manuscript has been restructured to improve clarity.

## References

F. Agostinis, C. Romualdi, G. Sales, and D. Risso. NewWave: a scalable R/Bioconductor package for the dimensionality reduction and batch effect removal of single-cell RNA-seq data. Bioinformatics, 38(9): 2648–2650, 2022.

J. Baglama and L. Reichel. Augmented implicitly restarted Lanczos bidiagonalization methods. SIAM Journal on Scientific Computing, 27(1):19–42, 2005.

K. Blighe, S. Rana, and M. Lewis. EnhancedVolcano: Publication-ready volcano plots with enhanced colouring and labeling, 2023. URL https://bioconductor.org/packages/EnhancedVolcano. R package version 1.18.0.

V. D. Blondel, J.-L. Guillaume, R. Lambiotte, and E. Lefebvre. Fast unfolding of communities in large networks. Journal of statistical mechanics: theory and experiment, 2008(10):P10008, 2008.

J. Cao, M. Spielmann, X. Qiu, X. Huang, D. M. Ibrahim, A. J. Hill, F. Zhang, S. Mundlos, L. Christiansen, F. J. Steemers, et al. The single-cell transcriptional landscape of mammalian organogenesis. Nature, 566 (7745):496–502, 2019.

T. Chari and L. Pachter. The specious art of single-cell genomics. PLOS Computational Biology, 19(8): e1011288, 2023.

V. Choulakian. Generalized bilinear models. Psychometrika, 61(2):271–283, 1996.

A. Duò, M. D. Robinson, and C. Soneson. A systematic performance evaluation of clustering methods for single-cell RNA-seq data. F1000Research, 7, 2018.

C. Eckart and G. Young. The approximation of one matrix by another of lower rank. Psychometrika, 1(3): 211–218, 1936.

I. N. Grabski, K. Street, and R. A. Irizarry. Significance analysis for clustering with single-cell RNA-sequencing data. bioRxiv, 2022.

C. Hafemeister and R. Satija. Normalization and variance stabilization of single-cell RNA-seq data using regularized negative binomial regression. Genome Biology, 20(1):1–15, 2019.

N. Halko, P.-G. Martinsson, and J. A. Tropp. Finding structure with randomness: Probabilistic algorithms for constructing approximate matrix decompositions. SIAM Review, 53(2):217–288, 2011.

V. Y. Kiselev, T. S. Andrews, and M. Hemberg. Challenges in unsupervised clustering of single-cell RNA-seq data. Nature Reviews Genetics, 20(5):273–282, 2019.

D. Lähnemann, J. Köster, E. Szczurek, D. J. McCarthy, S. C. Hicks, M. D. Robinson, C. A. Vallejos, K. R. Campbell, N. Beerenwinkel, A. Mahfouz, et al. Eleven grand challenges in single-cell data science. Genome Biology, 21(1):1–35, 2020.

J. Lause, P. Berens, and D. Kobak. Analytic Pearson residuals for normalization of single-cell RNA-seq UMI data. Genome Biology, 22(1):1–20, 2021.

J. T. Leek and J. D. Storey. Capturing heterogeneity in gene expression studies by surrogate variable analysis. PLoS Genetics, 3(9):e161, 2007.

F. Li, W. Won, E. Becker, J. Easlick, E. Tabengwa, R. Li, M. Shakhmatov, K. Honjo, P. Burrows, and R. Davis. Emerging roles for the fcrl family members in lymphocyte biology and disease. Fc Receptors, pages 29–50, 2014.

K. Z. Lin, J. Lei, and K. Roeder. Exponential-family embedding with application to cell developmental trajectories for single-cell rna-seq data. Journal of the American Statistical Association, 116(534):457–470, 2021.

R. Lopez, J. Regier, M. B. Cole, M. I. Jordan, and N. Yosef. Deep generative modeling for single-cell transcriptomics. Nature methods, 15(12):1053–1058, 2018.

M. D. Luecken and F. J. Theis. Current best practices in single-cell RNA-seq analysis: a tutorial. Molecular Systems Biology, 15(6):e8746, 2019.

A. Lun and M. Morgan. TENxBrainData: Data from the 10X 1.3 Million Brain Cell Study, 2020. R package version 1.8.0.

L. McInnes, J. Healy, and J. Melville. Umap: Uniform manifold approximation and projection for dimension reduction. arXiv preprint arXiv:1802.03426, 2018.

J. W. Miller and S. L. Carter. Inference in generalized bilinear models. arXiv preprint arXiv:2010.04896, 2020.

L. Morelli, V. Giansanti, and D. Cittaro. Nested stochastic block models applied to the analysis of single cell data. BMC Bioinformatics, 22(1):1–19, 2021.

Y. E. Nesterov. A method for solving the convex programming problem with convergence rate O(1/k2). In Dokl. Akad. Nauk SSSR,, volume 269, pages 543–547, 1983.

H. Pagès. DelayedArray: A unified framework for working transparently with on-disk and in-memory array-like datasets, 2020. R package version 0.14.1.

S. Petropoulos, D. Edsgärd, B. Reinius, Q. Deng, S. P. Panula, S. Codeluppi, A. P. Reyes, S. Linnarsson, R. Sandberg, and F. Lanner. Single-cell RNA-seq reveals lineage and X chromosome dynamics in human preimplantation embryos. Cell, 165(4):1012–1026, 2016.

H. A. Pliner, J. Shendure, and C. Trapnell. Supervised classification enables rapid annotation of cell atlases. Nature Methods, 16(10):983–986, 2019.

I. Razenshteyn, Z. Song, and D. P. Woodruff. Weighted low rank approximations with provable guarantees. In Proceedings of the Forty-Eighth Annual ACM Symposium on Theory of Computing, pages 250–263, 2016.

D. Risso, J. Ngai, T. P. Speed, and S. Dudoit. Normalization of RNA-seq data using factor analysis of control genes or samples. Nature Biotechnology, 32(9):896–902, 2014.

D. Risso, F. Perraudeau, S. Gribkova, S. Dudoit, and J.-P. Vert. A general and flexible method for signal extraction from single-cell RNA-seq data. Nature Communications, 9(1):1–17, 2018.

J. C. Roden, B. W. King, D. Trout, A. Mortazavi, B. J. Wold, and C. E. Hart. Mining gene expression data by interpreting principal components. BMC Bioinformatics, 7(1):1–22, 2006.

A.-E. Saliba, A. J. Westermann, S. A. Gorski, and J. Vogel. Single-cell RNA-seq: advances and future challenges. Nucleic Acids Research, 42(14):8845–8860, 2014.

A. Sarkar and M. Stephens. Separating measurement and expression models clarifies confusion in single-cell RNA sequencing analysis. Nature Genetics, 53(6):770–777, 2021.

R. Satija, J. A. Farrell, D. Gennert, A. F. Schier, and A. Regev. Spatial reconstruction of single-cell gene expression data. Nature Biotechnology, 33(5):495–502, 2015.

P. Savarese, D. McAllester, S. Babu, and M. Maire. Domain-independent dominance of adaptive methods. In Proceedings of the IEEE/CVF Conference on Computer Vision and Pattern Recognition, pages 16286–16295, 2021.

M. Soumillon, D. Cacchiarelli, S. Semrau, A. van Oudenaarden, and T. S. Mikkelsen. Characterization of directed differentiation by high-throughput single-cell RNA-seq. bioRxiv, page 003236, 2014.

N. Srebro and T. Jaakkola. Weighted low-rank approximations. In Proceedings of the 20th International Conference on Machine Learning (ICML), pages 720–727, 2003.

M. Stephens. False discovery rates: a new deal. Biostatistics, 18(2):275–294, 2017.

M. Stephens, P. Carbonetto, D. Gerard, M. Lu, L. Sun, J. Willwerscheid, and N. Xiao. ashr: Methods for Adaptive Shrinkage, using Empirical Bayes, 2022. URL https://CRAN.R-project.org/package=ashr. R package version 2. 2-54.

A. Subramanian, P. Tamayo, V. K. Mootha, S. Mukherjee, B. L. Ebert, M. A. Gillette, A. Paulovich, S. L. Pomeroy, T. R. Golub, E. S. Lander, et al. Gene set enrichment analysis: a knowledge-based approach for interpreting genome-wide expression profiles. Proceedings of the National Academy of Sciences, 102(43): 15545–15550, 2005.

Y. Sun, N. R. Zhang, and A. B. Owen. Multiple hypothesis testing adjusted for latent variables, with an application to the AGEMAP gene expression data. The Annals of Applied Statistics, 6(4):1664–1688, 2012.

V. Svensson. Droplet scRNA-seq is not zero-inflated. Nature Biotechnology, 38(2):147–150, 2020.

V. Svensson, A. Gayoso, N. Yosef, and L. Pachter. Interpretable factor models of single-cell rna-seq via variational autoencoders. Bioinformatics, 36(11):3418–3421, 2020.

F. W. Townes. Generalized principal component analysis. arXiv preprint arXiv:1907.02647, 2019.

F. W. Townes and K. Street. glmpca: Dimension Reduction of Non-Normally Distributed Data, 2020. URL https://CRAN.R-project.org/package=glmpca. R package version 0.2.0.

F. W. Townes, S. C. Hicks, M. J. Aryee, and R. A. Irizarry. Feature selection and dimension reduction for single-cell RNA-Seq based on a multinomial model. Genome Biology, 20(1):1–16, 2019.

E. Tuzhilina and T. Hastie. Weighted low rank matrix approximation and acceleration. arXiv preprint arXiv:2109.11057, 2021.

M. Uhlén, L. Fagerberg, B. M. Hallström, C. Lindskog, P. Oksvold, A. Mardinoglu, Å. Sivertsson, C. Kampf, E. Sjöstedt, A. Asplund, et al. Tissue-based map of the human proteome. Science, 347(6220):1260419, 2015.

C. A. Vallejos, D. Risso, A. Scialdone, S. Dudoit, and J. C. Marioni. Normalizing single-cell RNA sequencing data: challenges and opportunities. Nature Methods, 14(6):565–571, 2017.

L. Van der Maaten and G. Hinton. Visualizing data using t-SNE. Journal of Machine Learning Research, 9 (11), 2008.

F. A. Van Eeuwijk. Multiplicative interaction in generalized linear models. Biometrics, pages 1017–1032, 1995.

A. J. Wilk, A. Rustagi, N. Q. Zhao, J. Roque, G. J. Martínez-Colón, J. L. McKechnie, G. T. Ivison, T. Ranganath, R. Vergara, T. Hollis, et al. A single-cell atlas of the peripheral immune response in patients with severe COVID-19. Nature Medicine, 26(7):1070–1076, 2020.

S. Z. Wu, G. Al-Eryani, D. L. Roden, S. Junankar, K. Harvey, A. Andersson, A. Thennavan, C. Wang, J. R. Torpy, N. Bartonicek, et al. A single-cell and spatially resolved atlas of human breast cancers. Nature Genetics, 53(9):1334–1347, 2021.

G. X. Zheng, J. M. Terry, P. Belgrader, P. Ryvkin, Z. W. Bent, R. Wilson, S. B. Ziraldo, T. D. Wheeler, G. P. McDermott, J. Zhu, et al. Massively parallel digital transcriptional profiling of single cells. Nature Communications, 8(1):1–12, 2017.

H. Zou, T. Hastie, and R. Tibshirani. Sparse principal component analysis. Journal of Computational and Graphical Statistics, 15(2):265–286, 2006.

